# A mechanistic model for the negative binomial distribution of single-cell mRNA counts

**DOI:** 10.1101/657619

**Authors:** Lisa Amrhein, Kumar Harsha, Christiane Fuchs

## Abstract

Several tools analyze the outcome of single-cell RNA-seq experiments, and they often assume a probability distribution for the observed sequencing counts. It is an open question of which is the most appropriate discrete distribution, not only in terms of model estimation, but also regarding interpretability, complexity and biological plausibility of inherent assumptions. To address the question of interpretability, we investigate mechanistic transcription and degradation models underlying commonly used discrete probability distributions. Known bottom-up approaches infer steady-state probability distributions such as Poisson or Poisson-beta distributions from different underlying transcription-degradation models. By turning this procedure upside down, we show how to infer a corresponding biological model from a given probability distribution, here the negative binomial distribution. Realistic mechanistic models underlying this distributional assumption are unknown so far. Our results indicate that the negative binomial distribution arises as steady-state distribution from a mechanistic model that produces mRNA molecules in bursts. We empirically show that it provides a convenient trade-off between computational complexity and biological simplicity.

**Graphical Abstract:** 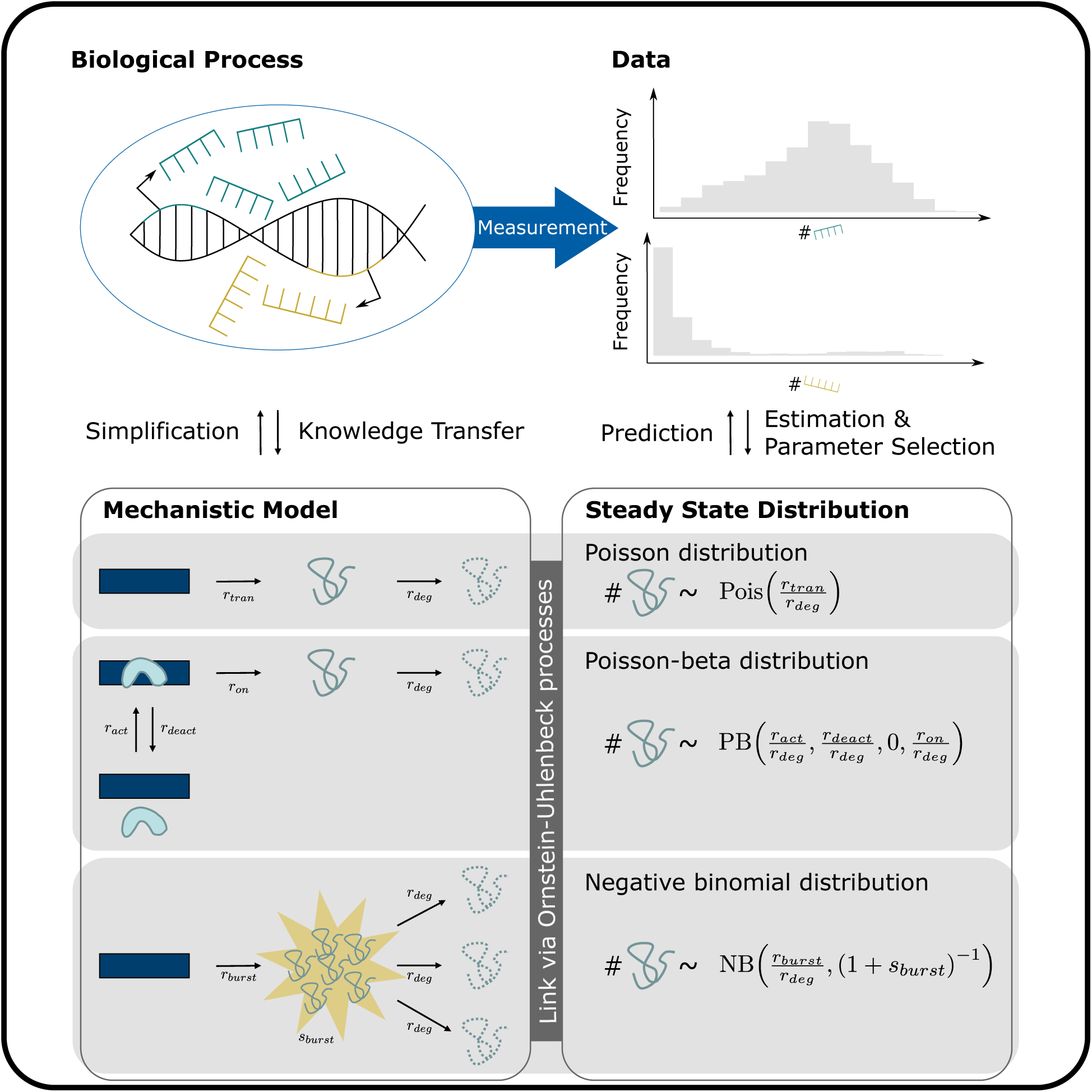

## Introduction

When analyzing the outcomes of single-cell RNA sequencing (scRNA-seq) experiments, it is essential to appropriately take properties of the resulting data into account. Many methods assume a parametric distribution for the sequencing counts due to its larger power than non-parametric approaches. To that end, a family of parametric distributions which adequately models the data needs to be chosen. In Supplementary Table S1, we provide an overview of computational tools for scRNA-seq analyses and their distribution choices. Among the 23 listed tools, around 60% use a negative binomial (NB) distribution, 40% a zero-inflated distribution (these two cases can overlap) and about 7% a Poisson-beta (PB) distribution.

Count data is most appropriately described by discrete distributions unless count numbers are without exception very high. A commonly chosen distribution is the Poisson distribution, which can be derived from a simple birth-death model of mRNA transcription and degradation. However, due to widespread overdispersed data, it is seldom suitable. Another typical choice is a three-parameter PB distribution (Delmans and Hemberg, 2016, Vu et al., 2016) which can be derived from a DNA switching model (also called *random telegraph model*, see Dattani and Barahona, 2017, or *basic model of gene activation and inactivation*, see Raj et al., 2006). Parameters of the PB distribution can be estimated from scRNA-seq data (Kim and Marioni, 2013), as well as experimentally measured and inferred (Suter et al., 2011). This distribution provides good estimates of scRNA-seq data; however, it entails the estimation of three parameters which introduces a high computational cost (Kim and Marioni, 2013). A frequent third choice is the NB distribution, used by several tools that analyze single-cell gene expression measurements such as SCDE (Kharchenko et al., 2014), Monocle 2 (Qiu et al., 2017) and many more (see Supplementary Table S1). This distribution is chosen due to computational convenience and good empirical fits. Mathematically, it can be considered as asymptotic steady-state distribution of the switching model (see Raj et al., 2006). However, this will entail biologically unrealistic assumptions. So far, no mechanistic model is known that directly leads to a NB distribution in steady state.

To close this gap, we look again at the already known mechanistic processes and their inferred parametric steady-state distributions: Poisson and PB. Integrating these in the general framework of Ornstein-Uhlenbeck (OU) processes (Barndorff-Nielsen and Shephard, 2001), we aim to transfer a general method of connecting mechanistic processes via stochastic differential equations (SDEs) and their theoretical steady-state distributions to this research problem. Hence, we show how to connect a desired steady-state distribution of the intensity process with the corresponding SDE by using OU processes and their properties. We use this method to calculate the corresponding SDE from the NB distribution as given steady-state distribution; from this, we can read a corresponding mechanistic model. In a (Case Study), we use our `R` package **scModels** to estimate three count distribution models (Poisson, PB and NB) on simulated perfect-world data, and perform model selection as well as goodness-of-fit tests. A comparison with existing implementations of the PB distribution, detailed derivations and definitions of the employed probability distributions can be found in the Appendix. Lastly, we repeat this comparison on real-world data and extend the models to more realistic ones by including zero inflation and heterogeneity.

By inferring a mechanistic model for stochastic gene expression, our work validates the NB distribution as a steady-state distribution for mRNA content in single cells.

## Results

It has previously been shown how to derive an mRNA count distribution from a simple birth-death model for mRNA transcription and degradation (Dattani and Barahona, 2017, Peccoud and Ycart, 1995). Alterations in the transcription and degradation model lead to alterations in the resulting mRNA count distribution. We will sketch the derivation of several such models and distributions. Our models describe the number of mRNA molecules in a cell for either *one* gene or for a group of genes for which we can assume identical kinetic parameters.

In the general context, we consider a transcription-degradation model with stochastic time-varying transcription rate *R*_*t*_ and deterministic constant degradation rate *r*_*deg*_ (Figure 1A). Here, the number of mRNA molecules at time *t* is Poisson distributed with intensity *I*_*t*_ following the random differential equation

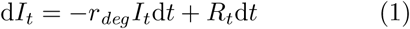

**Figure 1:**
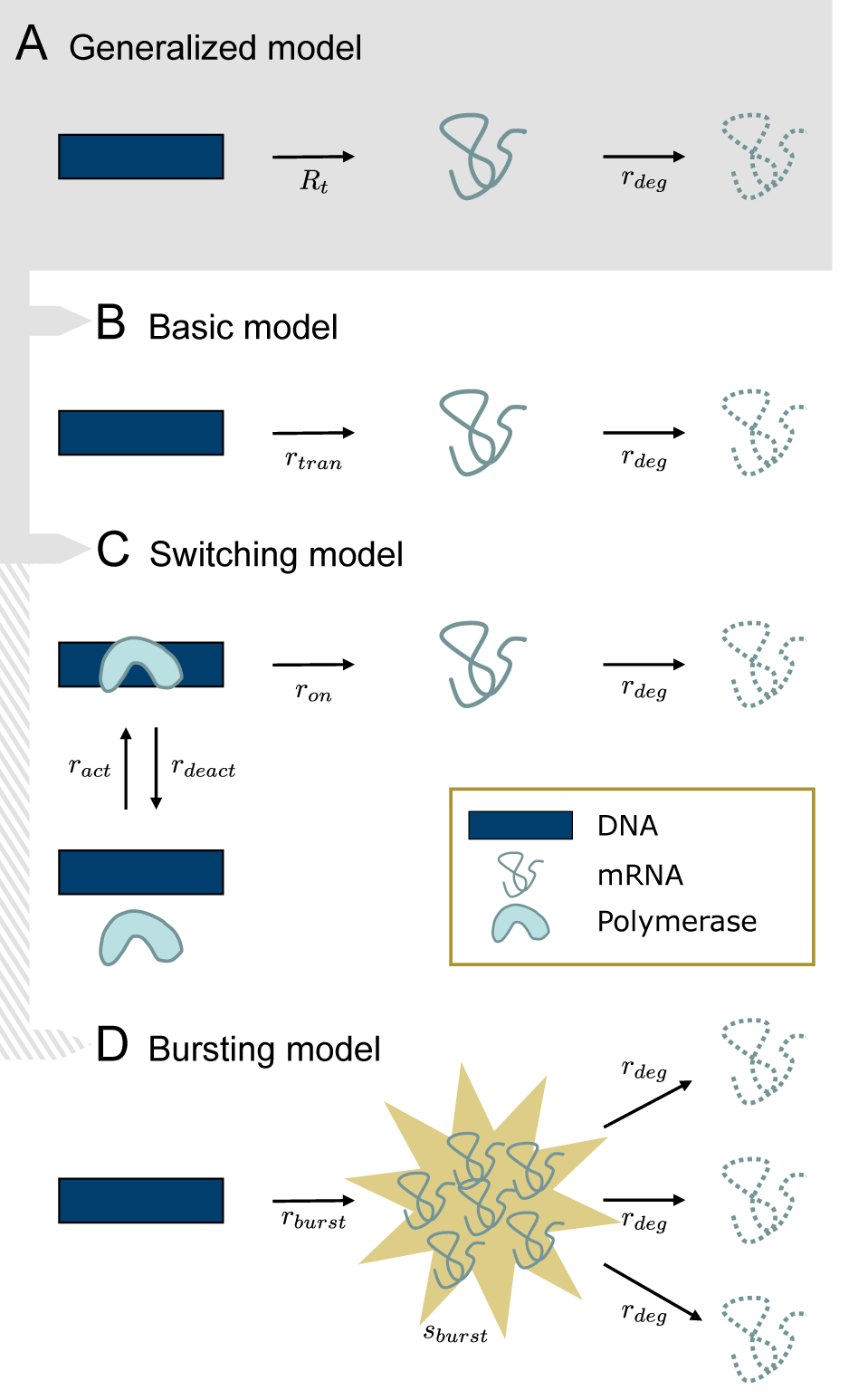
Transcription and degradation models: (A) Generalized model with time-dependent stochastic transcription rate *R*_*t*_ and constant deterministic degradation rate *r*_*deg*_. (B) Basic model with constant deterministic transcription and degradation rates. (C) Switching model of gene activation and inactivation, transcription and degradation. (D) Bursting model, where bursts occur at rate *r*_*burst*_ and burst sizes have mean *s*_*burst*_. This model differs from (A) in that transcription events can produce more than one mRNA molecule.

for *t* ≥ 0 and fixed *I*_0_ = *i*_0_ > 0. Depending on the transcription process, described by *R*_*t*_, this RDE has different solutions which will be shown in the following (for detailed calculations see Appendix).

### Basic model: constant transcription and degradation

In the *basic model*, transcription and degradation occur at constant rates *r*_*tran*_ and *r*_*deg*_ (Figure 1B). The RDE (1) simplifies to the ordinary differential equation (ODE)

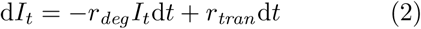

with time-independent non-stochastic steady state *I*_*t*_ = *r*_*tran*_ */r*_*deg*_. Hence, if the cell is in steady state, mRNA counts in this model follow a Poisson distribution with constant intensity *r*_*tran*_ */r*_*deg*_ (see Appendix).

### Switching model: gene activation and deactivation

In the well-known *switching model*, a gene switches between an inactive state where transcription is impossible, and an active state where transcription occurs. This can be explained by polymerases binding and unbinding to the specific gene (Figures 1C and S1). The RDE (1) becomes

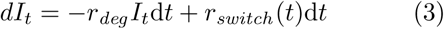

with

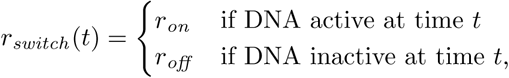

where *r*_*off*_ < *r*_*on*_. The transcription rate is modeled by a continuous-time Markov process (*r*_*switch*_ (*t*))_*t*≥0_ that switches between two discrete states *r*_*on*_ and *r*_*off*_ with activation and deactivation rates *r*_*act*_ and *r*_*deact*_, respectively. One usually sets *r*_*off*_ = 0. This corresponds to a system where a gene’s enhancer sites can be bound by different transcription factors or co-factors. Once bound, transcription occurs at constant rate *r*_*on*_, and mRNA continuously happens at constant rate *r*_*deg*_. Waiting times between switches are assumed to be exponentially distributed. As shown in the Appendix, these assumptions lead to *I*_*t*_ following a four-parameter Beta (*r*_*act*_ */r*_*deg*_, *r*_*deact*_ */r*_*deg*_, *r*_*off*_ */r*_*deg*_, *r*_*on*_ */r*_*deg*_) distribution, and therefore the mRNA content in steady state is described by a Poisson-beta (PB) distribution. Hence, the probability of having *n* mRNA molecules at time *t* is time-independent. For *r*_*off*_ = 0 (i. e. no transcription possible during inactive DNA state), it can be simplified to

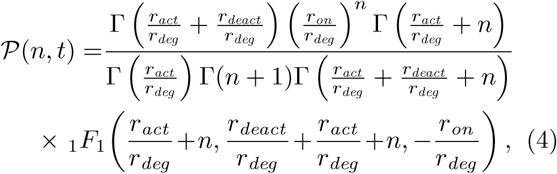

where Γ denotes the gamma function and 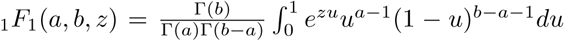 is the confluent hypergeometric function of first order, also called Kummer function. The density function of this PB distribution converges to the density function of a negative binomial (NB) distribution under specific conditions (Appendix). For *r*_*on*_ = *r*_*tran*_, *r*_*act*_ → ∞ and *r*_*deact*_ = 0, the switching model reduces to the basic model, and the above PB distribution collapses to a Poisson distribution with intensity parameter *r*_*tran*_ */r*_*deg*_, in consistency with the above-derived results.

### Connecting SDEs with steady-state distributions

Taken together, both models described the intensity process of a Poisson distribution (Equations 2 and 3). These intensity processes govern the transcription and degradation within the mechanistic models. They determine the steady-state distribution of the intensity parameter, and thus the overall distribution of the mRNA content. Importantly, changes in the intensity process lead to different steady-state distributions. We generalize this framework by using Ornstein-Uhlenbeck (OU) processes and their properties (Barndorff-Nielsen and Shephard, 2001).

The general definition of an OU SDE (adjusted to the above notation) is given by

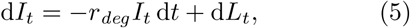

where *L*_*t*_ with *L*_0_ = 0 (almost surely) is a Lévy process, i. e. a stochastic process with independent and stationary increments. In addition, we need *L*_*t*_ to be a subordinator, that is a Lévy process with positive increments (Definition 7 in Appendix). A special property of OU processes is that, under certain conditions (see Definition 9), for a chosen distribution 𝒟 there is an OU process that in steady state leads to this distribution 𝒟. The other direction, i.e. the existence of a steady-state distribution 𝒟 for a chosen OU process (in terms of its subordinator), holds as well. For a given Lévy subordinator *L*_*t*_, the characteristic function of 𝒟, and thus 𝒟 itself, can be derived as described in the following (adjusted to the notation of Equation 5):

1. Find the characteristic function 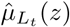 of the Lévy subordinator *L*_*t*_.
2. Calculate 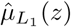 and write the result in the form exp(*ϕ*(*z*)) for some function *ϕ*(*z*).
3. Calculate the characteristic function *C*(*z*) of the stationary distribution 𝒟 of *I*_*t*_ by setting 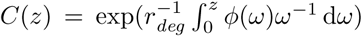. *C*(*z*) leads to 𝒟.

More details and examples are shown in the Appendix. Despite this apparently clear line of action, finding a corresponding law 𝒟 and process *L*_*t*_ is challenging without prior knowledge, e. g. if 𝒟 is not well-known or *L*_*t*_ is only specified through the characteristic function of *L*_1_. In the next section, we cast the NB distribution as an alternative distribution for which a subordinator can be derived.

### Negative binomial distribution: Deriving an explanatory bursting process

A widely considered model for scRNA-seq counts is the NB distribution. Like the above-employed PB distribution, it accounts for overdispersion by modeling the variance independently of the mean of the data. Having one parameter less than PB, NB is an appealing choice. However, mechanistic models underlying the NB distributional assumption are unknown. We aim to derive such a mechanistic model of transcription and degradation by reversing the steps that led from the switching model to the PB distribution. For that purpose, an important fact is that an NB distribution can be expressed as a Poisson-gamma (PG) distribution, i. e. as a conditional Poisson distribution with gamma distributed intensity parameter *I*. One has

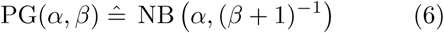

for *α, β* > 0 as derived in the Appendix.

In analogy to the derivation of the PB distribution from the switching model, we now seek to describe the mRNA content by a Poisson distribution with intensity parameter *I*_*t*_, which in steady state follows a gamma distribution instead of a beta distribution. Thus, we aim to specify an OU process with the gamma distribution as steady-state distribution. In terms of mechanistic modeling, this means that we need to describe a suitable transcription process. Mathematically, we need to specify the Lévy subordinator *L*_*t*_ accordingly. From financial mathematics it is known that a stationary gamma distribution is obtained if *L*_*t*_ is chosen to be a compound Poisson process (CPP, see Definition 8) with exponentially distributed jump sizes (Barndorff-Nielsen et al., 2001). This will be our choice of subordinator; however, the parameters of this process still need to be specified. In the following, we will show that the Lévy subordinator of the OU process (5) whose one-dimensional stationary distribution is Gamma(*α, β*), is a CPP with intensity parameter *α* · *r*_*deg*_ and mean jump size *β*^−1^.

To obtain this result, we follow the three-step procedure described above in reverse order. We start with 𝒟 ≙ Gamma(*α, β*) and transform its characteristic function to 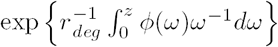, using the characteristic function of 𝒟 as given in the Appendix, Definition 1:

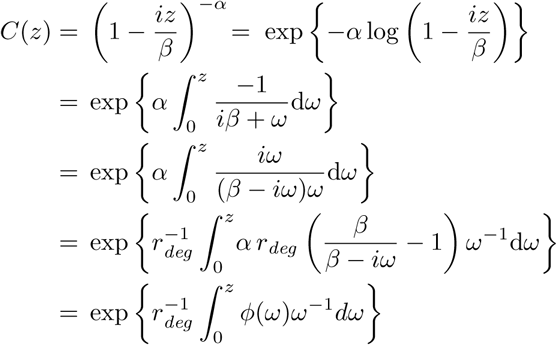

with 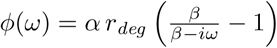 and *i* the imaginary number. Next, we use 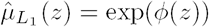 to obtain

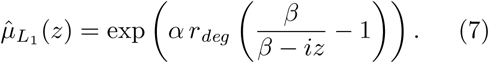

We aim to bring this into agreement with 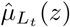, the time-dependent characteristic function of a general CPP *L*_*t*_ with intensity parameter *λ*. This is given by

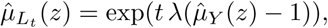

where *Y* is a random variable following the distribution of the jump sizes of the CPP, and 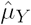 is its characteristic function (see Appendix, Definition 8). A CPP with intensity *λ* = *α* · *r*_*deg*_ and i.i.d. exponentially distributed increments *Y* ∼ Exp(*β*) with characteristic function 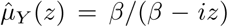 yields the overall characteristic function

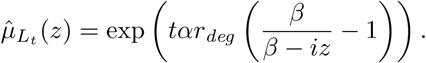

This is in accordance with 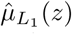 as derived in Equation (7), and hence, a mathematically appropriate subordinator is a CPP with intensity parameter *α* · *r*_*deg*_ and mean jump size *β*^−1^.

As a consequence, transcription is expressed via a stochastic process *L*_*t*_, namely the CPP, which experiences jumps after exponentially distributed waiting times. In contrast to the Lévy subordinators of the basic model, 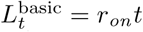, and of the switching model, 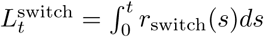, it possesses point-wise discontinuous sample paths (Figure 2, right). Intervals without any transcription activity seem to be disrupted by sudden explosions of mRNA numbers. This burstiness led us to call the mechanism behind the NB distribution the *bursting model*. We denote its subordinator by 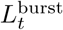 and argue the biological justification of the model in the Discussion and Conclusion.

**Figure 2:**
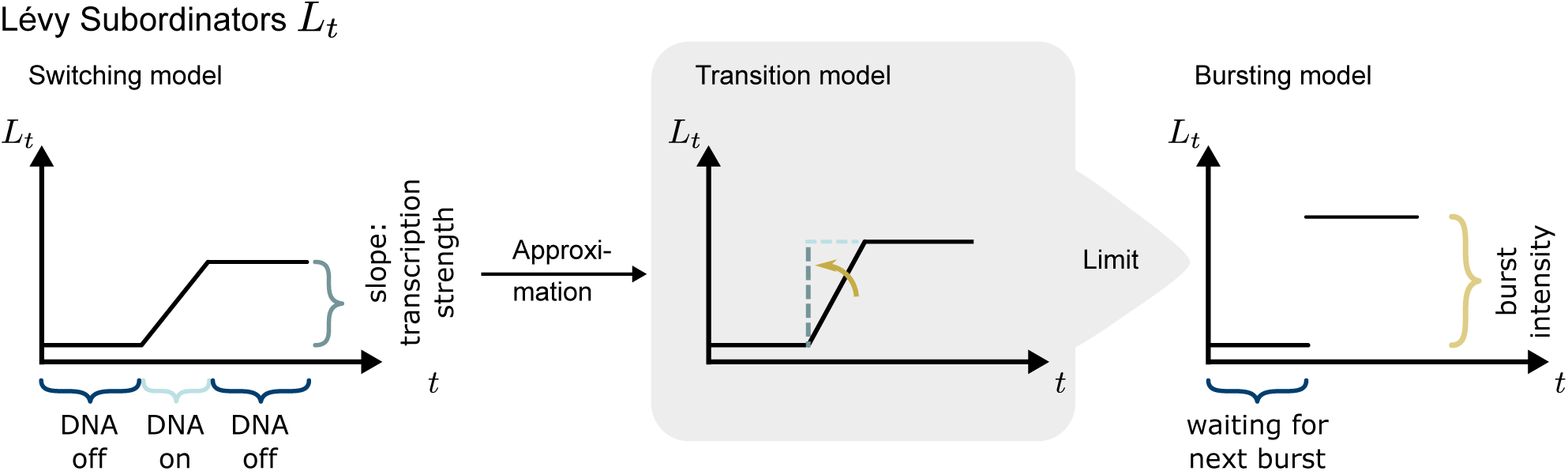
The Lévy subordinator of the switching model is shown on the left by means of an exemplary trajectory. For small duration of the DNA being active and large transcription strength, its behavior can be approximately described by a step function as depicted in the middle (transition model). The limit of this approximation, with infinitesimally small DNA activation time interval and infinitesimally large transcription strength, leads to a trajectory of the subordinator of the bursting model which is shown on the right.

We aim to derive the mechanistic transcription process of the bursting model in more detail. Specifically, we tackle the distribution of burst sizes of mRNA counts. For this we look at a heuristic transition from 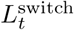 to 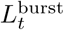.

First, we dismantle the shape of the trajectories of 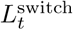. As depicted in Figure 2 on the left, such a trajectory consists of alternating piecewise constant and piecewise linear parts. The constant parts appear during time intervals without transcription, i. e. where the DNA is inactive. The length of such a time interval depends only on the rate *r*_*act*_ of the switching model. Once the DNA switches into the active mRNA transcribing state, the time interval with transcription depends only on the rate *r*_*deact*_. The slope of the trajectory during this active DNA state represents the transcription strength and depends only on the parameter *r*_*on*_.

In case the length of the time interval of active DNA becomes infinitesimally small, and at the same time the transcription strength becomes infinitesimally large, the trajectory of 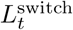 turns into a step function as depicted in Figure 2 on the right. This limit is obtained if *r*_*deact*_ → ∞ and *r*_*on*_ → ∞ in a way that needs to be specified. For that reason, we in the following seek to describe a mechanistic model for the transition phase (Figure 2, middle) leading to the bursting model.

In the switching model, as soon as DNA becomes active, a competition starts between the events *transcription* and *deactivation*. In addition, degradation may happen, which will affect the intensity process *I*_*t*_ and the number of mRNA molecules, but not the transcription process. If a transcription event occurs, the competition between transcription and deactivation continues at the same probability rates as before; the only affected event probability is the one for degradation because this probability depends on the current mRNA count. We now consider the following approximation of the switching model and call it the *transition model* : When DNA becomes active, we allow the events transcription and deactivation to happen, but not degradation. To correct for the missing degradation events, we introduce a waiting time *W* after DNA deactivation in which only degradation can occur, but no DNA activation. For appropriately chosen *r*_*deact*_ → ∞ and *r*_*on*_ → ∞, the approximation error will tend to zero.

The number of transcription events *S* during one active DNA phase is geometrically distributed with success probability parameter *r*_*deact*_ */*(*r*_*deact*_ + *r*_*on*_). In the interpretation of the geometric distribution, transcription events are considered as failures, deactivation as success. The waiting time *W* needs to accumulate the times before *S* transcriptions and one deactivation. Thus, *W* = *T*_1_ + … + *T*_*S*_ + *D*, where *T*_*i*_ ∼ Exp(*r*_*on*_), *i* = 1, …, *S*, are the single waiting times for each transcription event and *D* ∼ Exp(*r*_*deact*_) is the waiting time till the next DNA deactivation.

Taken together, the bursting process can be considered as the limiting process of the approximation of the switching process as *r*_*on*_ → ∞ and *r*_*deact*_ → ∞ under the condition that the success probability parameter of the geometric distribution, *r*_*deact*_ */*(*r*_*deact*_ + *r*_*on*_) stays constant. As the link between the switching model and PB distribution is known, and since PB converges towards NB under certain conditions (see Appendix), we can connect the parameters of the bursting model with those of the NB distribution and CPP.

That is, the bursting model can mechanistically be described as follows: After Exp(*r*_*burst*_)-distributed waiting times, a Geo((1 + *s*_*burst*_)^−1^)-distributed number of mRNAs are produced at once, where *s*_*burst*_ is the mean burst size (see also Golding et al., 2005). As in the basic and switching models, degradation events occur with Exp(*r*_*deg*_)- distributed waiting times. The just described mechanistic bursting model is shown in Figure 1D. It can equivalently be described by the OU process (5) with *L*_*t*_ being a CPP with Exp(*r*_*burst*_)-distributed waiting times and Exp(*s*_*burst*_)-distributed jump sizes. Thus, in steady state, mRNA counts follow a PG(*r*_*burst*_ */r*_*deg*_, *s*_*burst*_) distribution or, equivalently, a NB (*r*_*burst*_ */r*_*deg*_, (1 + *s*_*burst*_)^−1^) distribution if the bursting model is assumed.

The NB (*r*_*burst*_ */r*_*deg*_, (1 + *s*_*burst*_)^−1^) model, again, can be interpreted as follows (see also Appendix, Definition 3): Assume you have an empty bucket into which you put balls according to the following stochastic procedure. You perform a number of independent Bernoulli trials, each with success probability (1 + *s*_*burst*_)^−1^. If there is a failure, you add one ball to the bucket. If there is a success, you do not do anything but count the success event. You continue until there have been *r*_*burst*_ */r*_*deg*_ successes. (For interpretation purposes, we here assume *r*_*burst*_ */r*_*deg*_ to be a whole-valued number.) The larger *s*_*burst*_, the smaller the success probability, i. e. by expectation you will put more balls in the bucket for large *s*_*burst*_. Similarly, the larger the ratio of *r*_*burst*_ to *r*_*deg*_, the more success events will be waited for, thus the more balls will tend to be added. The final number of balls in the bucket represents the number of mRNA molecules in a cell at steady state.

The above top-down derivation from the steady-state distribution to the mechanistic process has to be motivated heuristically in parts. In the Appendix we prove bottom-up that the above described mechanistic bursting model indeed leads to the steady-state NB distribution by directly calculating the master equation (see also Supplementary Figure S2).

### Heterogeneity and dropout

The transcription and degradation models considered so far describe the number of mRNA molecules for homogeneously expressed genes that are actually present in a cell. Real-world data is usually more complex: First, cell populations may be heterogeneous. Second, scRNA-seq measurements will be subject to measurement error. For example, they often contain a large number of zeros. If a zero is due to technical error, it is called dropout. Regardless of what causes this phenomenon, a data model should take this property into account. We describe two model extensions here.

Data that originates from different cell populations (in terms of different transcriptomic properties) can be modeled mathmatically. If a population consists of e. g. two subpopulations, each of them is modeled by one single distribution, 𝒟_1_ or 𝒟_2_, parameterized via *θ*_1_ and *θ*_2_, respectively. One assumes the mRNA count to be distributed according to *p*𝒟_1_(*θ*_1_)+(1 − *p*)𝒟_2_(*θ*_2_) with *p* ∈ [0, 1], that means: With probability *p*, the count distribution of that cell is 𝒟_1_, otherwise 𝒟_2_. The corresponding mixture density is given by

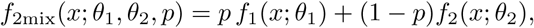

where *f*_1_ and *f*_2_ are the densities of 𝒟_1_ and 𝒟_2_, and *x* is the observed mRNA count. For *k* > 2 subpopulations, the density can easily be generalized to a mixture of *k* distributions 𝒟_1_, …, 𝒟_*k*_ with probabilities *p*_1_, …, *p*_*k*_:

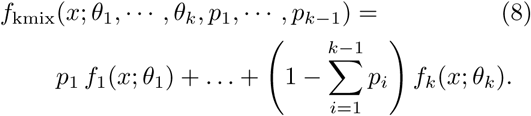

The distributions 𝒟_*i*_ can be any (ideally discrete count) distribution, possibly from different distribution families.

An appropriate model for the occurrence of the above-mentioned dropout is a zero-inflated distribution (Kharchenko et al., 2014). For one homogeneous population, the mRNA count will be distributed according to *p*𝟙_{0*}*_ +(1 − *p*)𝒟(*θ*), with 𝟙_{0}_ being the indicator function with point mass at zero, and the corresponding density function reads

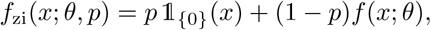

where *f* is the density function of 𝒟. Analogously, zero inflation can be added to a mixture of several distributions, see (8). mRNA counts are then distributed according to

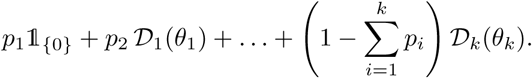

The corresponding density function is given by

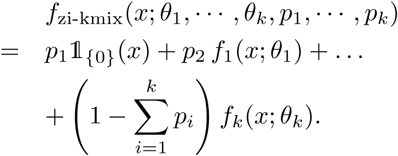

### Data application

We perform a comprehensive comparison of the considered mRNA count distributions, that is the Poisson, NB and PB distribution, when applied to real-world data. Within each of the three distributions we further consider mixtures of two populations (from identical distribution types but with different parameters) with and without additional zero inflation. In total, we investigate twelve different models as shown in Figure 3. The numbers of parameters in these models are listed in Supplementary Table S3.

**Figure 3:**
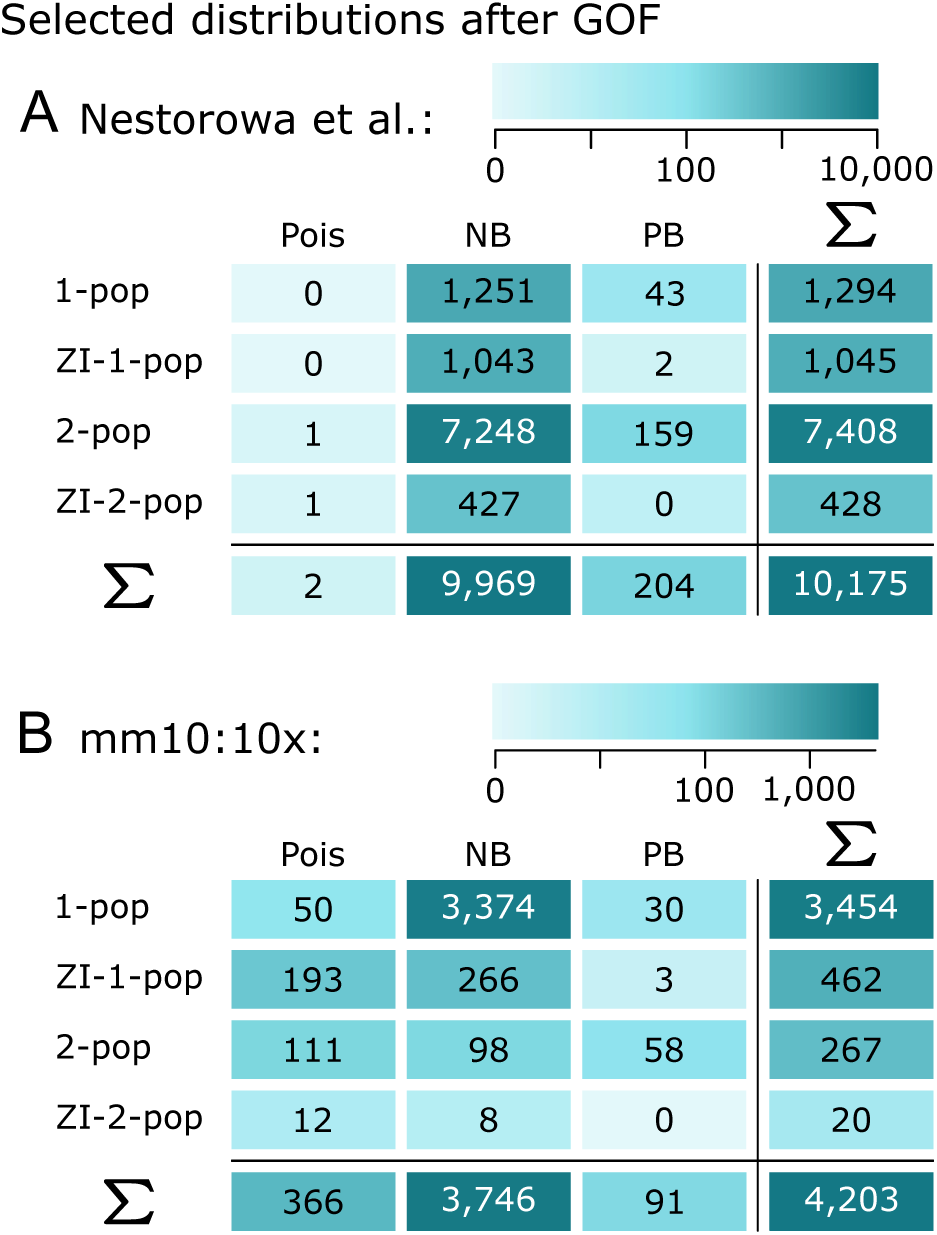
Frequencies of chosen distributions via BIC-after-GOF applied to real-world datasets: (A) Nestorowa et al. (2016) (B) mm10:10x, Official 10x Genomics Support (2017).

We estimate these twelve models on two real-world datasets: The first one stems from Nestorowa et al. (2016), contains 1,656 mouse HSPCs and was generated using the Smart-Seq2 (Picelli et al., 2014) protocol, and thus did not employ unique molecular identifiers (UMIs). The second dataset contains 3,356 homogeneous NIH3T3 mouse cells and has been generated using the 10x Chromium technique (Zheng et al., 2017), thus incorporating UMIs. It is part of the publicly available 10x dataset “6k 1:1 Mixture of Fresh Frozen Human (HEK293T) and Mouse (NIH3T3) Cells” (Official 10x Genomics Support, 2017). Here, we refer to this dataset as *mm10:10x*.

We applied a gene filter (see Appendix), estimated the model parameters of the twelve considered models via maximum likelihood, and performed model selection as described in the Case Study via the Bayesian information criterion (BIC). Figure 3 summarizes the frequencies of the chosen models across genes. We only display those choices where the chosen distribution with estimated parameters was not rejected by a goodness-of-fit (GOF) test (*χ*^2^-test) at 5% significance level with multiple testing correction.

In the data from Nestorowa et al. (2016), 16,364 genes remained after filtering, of which 10,175 were not rejected by the GOF test. Figure 3A shows that some variant of the NB distribution was chosen for 98% of these genes. Among these, mRNA count numbers for many genes were best described by the zero-inflated NB distribution. However, an even higher preference could be observed for the mixture of two NB distributions. This can be explained by taking a closer look at the gene expression counts of the affected genes (see also Supplementary Figure S6): Most of those genes not only show many zeros, but also many low non-zero counts, i. e. many ones, twos etc., next to higher counts. Such expression profiles are not covered by a simple zero-inflated model but prefer a mix of two distributions, one of them mapping to low expression values (see Supplementary Table S3).

In the mm10:10x data, 4,203 genes remained after filtering and the GOF test. Figure 3B shows that for 89% of these genes, an NB distribution variant was chosen as most appropriate model. However, other than for the dataset from Nestorowa et al. (2016), the standard NB distribution (for one population, without zero inflation) was sufficient in the majority of cases. We looked for commonalities between the gene profiles that led to the same distribution choice. Supplementary Figure S5 suggests an interdependence between the chosen one-population distributions and the range of the parameter estimates.

Taken together, the NB distribution was chosen for most gene profiles, either as a single distribution, a mixture of two NB distributions or with additional zero inflation.

### NB distribution as commonly chosen count model

While the mechanistic models and their steady-state distributions describe actual mRNA contents in single cells, real-world data underlies technical variation in addition to biological complexity. We investigated in a simulation study (Case Study and Figure 4) and on real-world data (Figure 3) which distributions were most appropriate among those considered to describe gene expression profiles. The simulation study showed that an NB distribution may be best suited even if the *in silico* data had been generated from the switching model. Also in the real-data application, the NB distribution was chosen in most cases. In line with our expectations, gene profiles of the non-UMI-based dataset by Nestorowa et al. (2016) showed strong preference for a two-population mixture or zero-inflated variant of the NB distribution. In contrast, the mm10:10x dataset consists by construction of homogeneous cells, and 10x Chromium is not known for large amounts of unexpected zeros in the measurements. Accordingly, the single-population NB distribution was sufficient for most gene profiles here. For 9% of the considered genes in the m10:10x dataset, mRNA counts were most appropriately described by some form of the Poisson distribution. We have examined these 366 genes for functional similarities; while estimated parameters show some apparent pattern (Supplementary Figure S5), we did not find any defining biological characteristics (Supplementary Figure S7).

**Figure 4:**
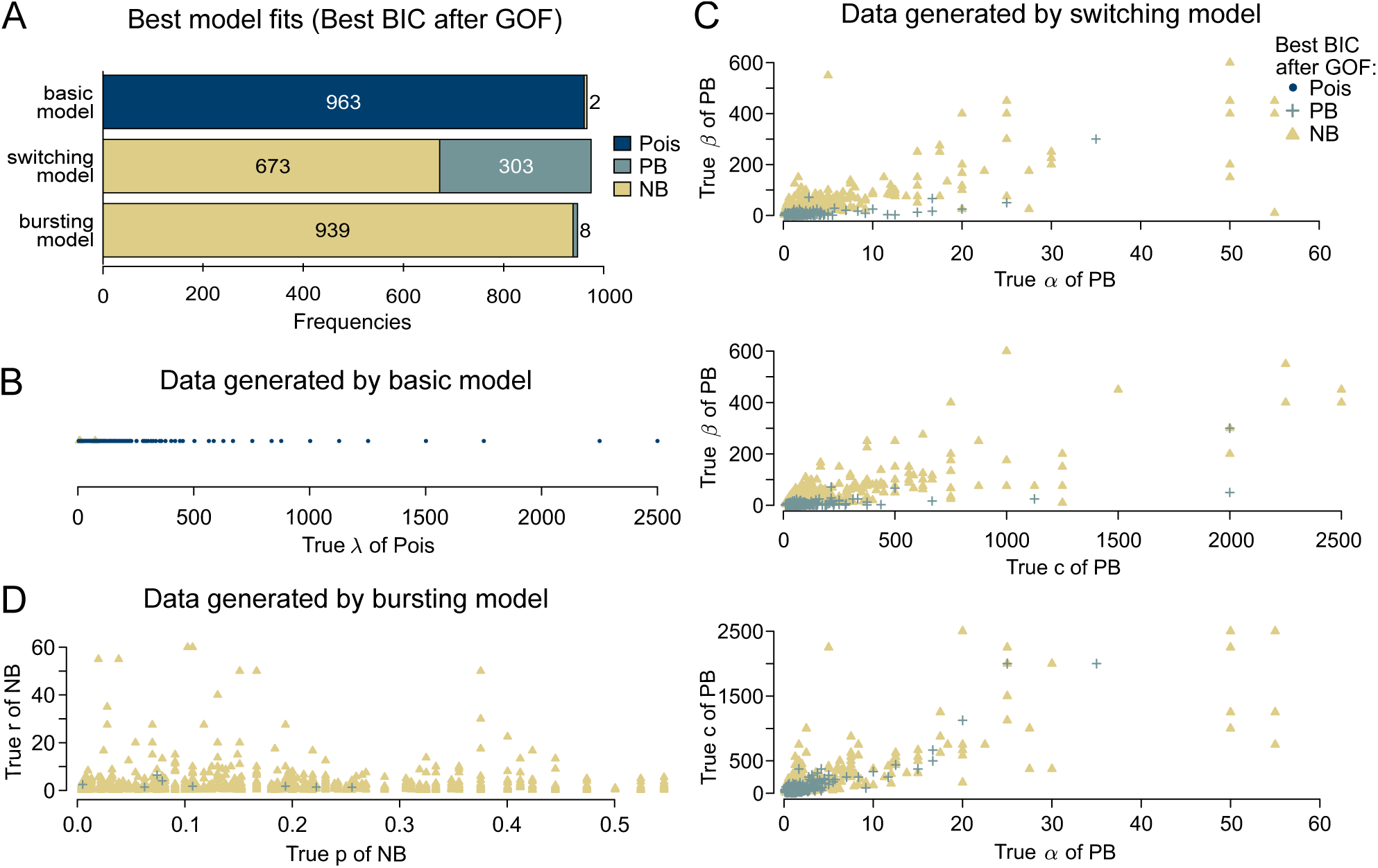
Model selection on *in silico* data: (A) Frequencies of chosen distributions (Poisson, PB, NB) via BIC-after-GOF based on datasets generated by the three different transcription models (basic, bursting, switching). (B-D) Employed parameter values (indicated by horizontal/vertical position) and chosen distributions (indicated by color/symbol) for basic model (B), switching model (C) and bursting model (D). The names of the parameters correspond to those in Definitions 2, 3 and 5 in the Appendix.

Similar to us, Vieth et al. (2017) performed model selection among Poisson, NB and PB distributions by BIC and GOF on several publicly available datasets. Although they used the method of Vu et al. (2016) for which computation of the GOF statistics is impossible for the PB distribution, they still observed a tendency towards the NB distribution as preferred models. In our study, we represent the PB density in terms of the Kummer function, which allows us to compute the GOF statistics accordingly.

Different sequencing protocols might lead to differences in distributions and also might generate data of different magnitudes. Ziegenhain et al. (2017) applied various sequencing methods to cells of the same kind to understand the impact of the experimental technique on the data. Based on the so-generated data, Chen et al. (2018) investigated differences in gene expression profiles between read-based and UMI-based sequencing technologies. They concluded that, other than for read counts, the NB distribution adequately models UMI counts. Townes et al. (2019) suggest to describe UMI counts by multinomial distributions to reflect the nature of the sequencing procedure; for computational reasons, they propose to approximate the multinomial density again by an NB density. Overall, the NB distribution appears sufficiently flexible to hold independently of the specific sequencing approach.

### R package scModels

We provide the `R` package **scModels** which contains all functions needed for maximum likelihood estimation of the considered distribution models. Three applications of the Gillespie algorithm (Gillespie, 1976) allow synthetic data simulation (as used in the Case Study) via the basic, switching and bursting model, respectively. Implementations of the likelihood functions for the one-population case and two-population mixtures, with and without zero inflation, allow inference of the Poisson, NB and PB distributions. We provide a new implementation of the PB density, based on our novel implementation of the Kummer function, also known as the generalized hypergeometric series of Kummer. This became necessary, because the existing `R` function (kummerM() contained in package **fAsianOptions**) was only valid for specific parameter values, and hence, was not suited for optimization in continuous unconstrained space (more information in Appendix). Existing packages such as **D3E** (Delmans and Hemberg, 2016), implemented in `Python`, and **BPSC** (Vu et al., 2016), implemented in `R`, use the PB distribution for scRNA-seq data analysis but based on a different approximation scheme (see Appendix). With our new implementation of the PB density we did not overcome the problem of time-consuming calculation, but we for the first time provided an implementation of the Kummer function in `R` valid for all values required inside the PB density. For a more detailed description of **scModels** and a package comparison to **D3E** and **BPSC**, see the Appendix.

## Case Study

In a simulation study, we generate *in silico* data from the three considered mechanistic models: the basic model (Figure 1B), the switching model (Figure 1C), and the bursting model (Figure 1D), using the Gillespie algorithm implemented in **scModels**. In order to choose realistic values for the rate parameters, we orient ourselves on experimental studies which aim to determine rates of the switching process in specific cases. For example, Suter et al. (2011) identify rates for so-called short-lived genes where mRNA and protein pulses are directly connected to one single on-and-off-switch of a gene. We provide an overview of the employed rate combinations in Supplementary Table S4.

As a proof of concept, we estimate the three corresponding distributions, i. e. the Poisson, the Poisson-beta (PB) and the negative binomial (NB) distribution, on all generated datasets via maximum likelihood estimation. To investigate which distribution explains the data best, we compute the Bayesian information criterion

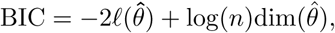

where *ℓ* represents the corresponding log-likelihood function, *θ* the possibly multivariable parameter vector of the distribution, 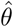 its maximum likelihood estimate, 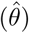 its dimension, i. e. the number of unknown scalar parameters, and *n* the sample size. The distribution with lowest BIC value is considered most appropriate among all considered models. Afterwards, we apply a *χ*^2^-test to assess the goodness-of-fit (GOF) and neglect those datasets for which the distribution fits are rejected at the 5% significance level (without multiple testing correction). This reduces the total number of 1,000 simulated datasets per model to the amounts displayed in Figure 4A.

We investigate whether the selected distributions correspond to the distributions that arise from the respective mechanistic models: For the datasets generated from the basic model, model selection via BIC-after-GOF indeed prefers the Poisson distribution in most cases, independently of the used distribution parameter *λ* (Figure 4A, first bar, and Figure 4B). In contrast, for datasets generated by the switching model, BIC-after-GOF in big parts chooses either the NB or the PB distribution (Figure 4A, middle bar). The choice seems to depend on the employed rate parameters: Figure 4C indicates a tendency towards the PB distribution for low values of *β*; otherwise, the NB distribution often seems to model the data generated by the switching model sufficiently well. For datasets generated by the bursting model, BIC-after-GOF picks the NB distribution for the majority of the time without any obvious bias (Figure 4A/D). The study shows that, apparently, the NB distribution is complex enough to describe the data generated from the switching model. The BIC decides in many cases that a potentially better fit is not worth the extra effort for estimating an additional parameter in the PB distribution.

## Discussion and Conclusion

In this work, we derived a mechanistic model for stochastic gene expression that results in the NB distribution as steady-state distribution for mRNA content in single cells. According to the so-obtained bursting model, transcription happens in chunks, rather than in a one-by-one production as commonly assumed in mechanistic modeling (Dattani and Barahona, 2017). We discuss the biological plausibility of bursty transcription further below. The consideration of the bursting model and its derivation is interesting from both practical and theoretical points of view:

First of all, the NB distribution is defined through two parameters whereas the PB distribution typically requires three parameters to be specified in the current context. Therefore, the NB distribution is computationally less elaborate to estimate, given some data, than the PB distribution. Several tools employ the NB distribution to parameterize mRNA read counts (see Supplementary Table S1). However, there has been no mechanistic biological model known so far leading to this distribution, other than for the Poisson and the PB distributions (Figures 1B,C). Here, we provide a possible explanation.

Second, we demonstrated how to generally link a probability distribution to an Ornstein-Uhlenbeck (OU) process and derive a mechanistic model. This brings a new field of mathematics to single-cell biology. The procedure can be used to deduce possible mechanistic processes leading to different steady-state distributions, exploiting the rich literature on OU processes from financial mathematics.

Third, although we focused on the resulting steady-state distributions of the mechanistic models here, our mathematical framework also provides model descriptions in terms of stochastic processes. Nowadays, sequencing counts are commonly available as snapshot data. However, time-resolved measurements may become standard (Golding et al., 2005), and in that case our models open up the statistical toolbox of stochastic processes to extract information from interdependencies within single-cell time series.

### Limiting cases of the switching model that give rise to the NB distribution are biologically unrealistic

The NB and PB distributions have been linked before. Among others, Raj et al. (2006) and Grün et al. (2014) have shown that the NB distribution is an asymptotic result of the switching model and the corresponding PB distribution (see Appendix). However, this result holds only under biologically unrealistic assumptions as we elaborate in the following. Our derivation of the NB steady-state distribution, in contrast, is based on a thoroughly realistic mechanism of bursty transcription. The approach by Raj et al. (2006) and Grün et al. (2014) requires *r*_*deact*_ */r*_*deg*_ → ∞ and *r*_*on*_ */r*_*deact*_ < 1. That means, the deactivation rate has to be substantially larger than the mRNA degradation rate and, simultaneously, the transcription rate needs to be smaller than the gene deactivation rate. Here, we discuss the plausibility of these presumptions:

Schwanhäusser et al. (2011) showed that mRNA half-life is in median around *t*_1*/*2_ = 9 *h* (range: 1.61 *h* to 40.47 *h*), which results in a degradation rate *r*_*deg*_ = log(2)*/t*_1*/*2_ of 0.077 *h*^−1^ = 0.00128 min^−1^ (range: 0.00718 min^−1^ to 0.00029 min^−1^). For *r*_*deact*_ */r*_*deg*_ → ∞, the mRNA degradation rate needs to become much smaller than the gene deactivation rate. Visual comparison shows that density curves of the PB and according NB distributions start to look similar for *r*_*deact*_ */r*_*deg*_ ≈ 20, 000. Assuming a 20,000-fold larger gene deactivation rate results in *r*_*deact*_ = 29.67 min^−1^ (range: 143.51 min^−1^ to 5.71 min^−1^). This means that on average the gene switches approximately 30 times per minute into the off-state, i. e. on average the gene is in its active state for only two seconds. RNA polymerases proceed at 30 nt/sec (without pausing at approximately 70 nt/sec) (Darzacq et al., 2007). Genes have a length of hundreds to thousands of nucleotides. Thus, according to the above numbers, genes cannot be transcribed in such short phases. The DNA needs to stay active during the whole transcription process of one (or more) mRNAs; as soon as the DNA turns inactive, all currently running transcriptions are stopped. In other words, although the NB distribution can mathematically be derived as a limiting steady-state distribution of the switching model, this entails biologically implausible assumptions.

This criticism is underpinned by the work of Suter et al. (2011) who derived ranges of the rates of the switching model experimentally and by calculations. Here, only so-called short-lived genes were taken into account. Thus, observed mRNA half-lives were on a smaller scale, mainly between 30 and 140 min, resulting in mRNA degradation rates between 0.005 min^−1^ and 0.023 min^−1^. At the same time, deactivation rates were found in the range between 0.1 min^−1^ and 0.6 min^−1^. Hence, their quotient is at maximum around 120 and thus nowhere close to infinity. Another mathematical assumption for deriving the NB limit distribution was that the transcription rate needed to be smaller than the deactivation rate. This is not confirmed by Suter et al. (2011) for most genes.

### Biological plausibility of bursting model

Burst-like transcription has been discussed, e. g. Golding et al. (2005), Schwanhäusser et al. (2011) and Suter et al. (2011). We take a look at the inherent assumptions of the bursting model: The bursting rate *r*_*burst*_ represents the waiting time until the DNA turns open for transcription in addition to the time which the polymerase needs to transcribe. The model assumes that several polymerases attach simultaneously to the DNA and terminate transcription at the same time. By simplifying this part of the transcription process model, the problem of persisting DNA activation during the whole transcription process in the switching model is avoided.

### Practical relevance

There is no unambiguous answer to the question of the most appropriate probability distribution for mRNA count data. Pragmatic reasons will often lead to NB distribution as already employed by many tools (see Supplementary Table S1). However, the choice may depend on experimental techniques, the statistical analysis to be performed, and also differ between genes within the same dataset. For large read counts, even continuous distributions may be most suitable.

While statistics quantifies which model is the most plausible one from the data point of view, mathematical modelling points out which biological assumptions may implicitly be made when a particular distribution is used. Importantly, while the mechanistic model leads to a unique steady-state distribution, the reverse conclusion is not true. In general, the basic model and the corresponding Poisson distribution may appear too simple in most cases (both with respect to biological plausibility and the ability to describe measured sequencing data). The switching and bursting models are harder to distinguish. From the mathematical point of view, their densities are of similar shape, such that the less complex NB model will often be preferred. Answering the question from the biological perspective may require measuring mRNA generation at a sufficiently small time resolution (e. g. Golding et al., 2005) to see whether several mRNA molecules are generated at once (bursting model) or in short successional intervals (switching model). Taken together, we have identified mechanistic models for mRNA transcription and degradation with good interpretability, and established a link to mathematical representations by stochastic processes and steady-state count distributions. Specifically, the commonly used NB model is supplied with a proper mechanistic model of the underlying biological process. The `R` package **scModels** over-comes a previous shortcoming in the implementation of the PB density. It provides a full toolbox for data simulation and parameter estimation, equipping users with the freedom to choose their models based on content-related, design-based or purely pragmatic motives.

## Supplemental Information

Supplemental Information includes seven figures and four tables which can be found at the end of this paper.

## Author Contributions

The study was designed by LA and CF. LA developed and performed the mathematical analysis and software development with help of KH and CF. LA and CF wrote the paper.

## Acknowledgments

Our research was supported by the German Research Foundation within the SFB 1243, Subproject A17, by the German Federal Ministry of Education and Research under grant number 01DH17024, and by the National Institutes of Health under grant number U01-CA215794.

## Appendix

### OVERVIEW TOOLS TABLE

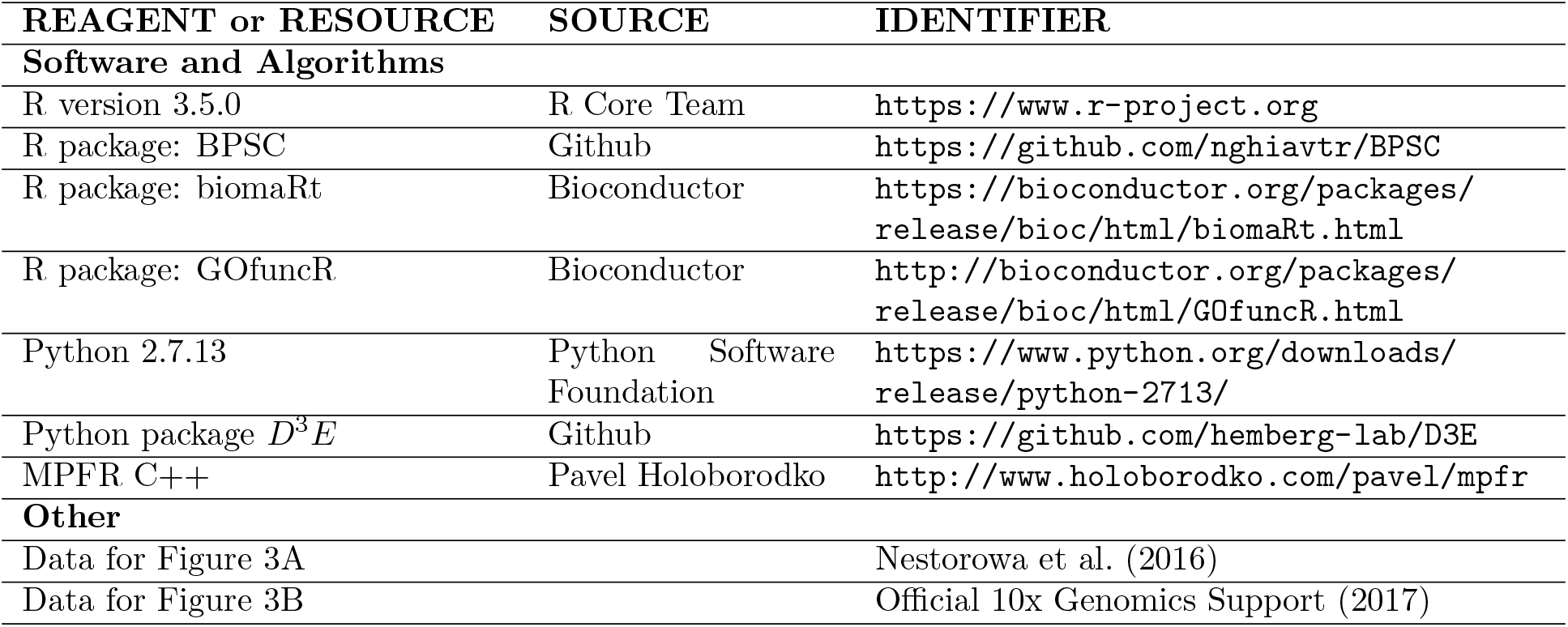

| REAGENT or RESOURCE | SOURCE | IDENTIFIER |
| --- | --- | --- |
| **Software and Algorithms** |  |  |
| R version 3.5.0 | R Core Team | https://www.r-project.org |
| R package: BPSC | Github | https://github.com/nghiavtr/BPSC |
| R package: biomaRt | Bioconductor | https://bioconductor.org/packages/release/bioc/html/biomaRt.html |
| R package: GOfuncR | Bioconductor | http://bioconductor.org/packages/release/bioc/html/GOfuncR.html |
| Python 2.7.13 | Python Software Foundation | https://www.python.org/downloads/release/python-2713/ |
| Python package $D^3E$ | Github | https://github.com/hemberg-lab/D3E |
| MPFR C++ | Pavel Holoborodko | http://www.holoborodko.com/pavel/mpfr |
| **Other** |  |  |
| Data for Figure 3A |  | Nestorowa et al. (2016) |
| Data for Figure 3B |  | Official 10x Genomics Support (2017) |

### METHOD DETAILS

#### DEFINITIONS AND IDENTITIES

Probability distributions and other mathematical terms are often not uniformly defined in literature. In this section, we explain the terminology used in the present work. References include Dormann (2013), the NIST library (Olver et al., 2019), Karlis and Xekalaki (2005), Rogers and Williams (2000), Barndorff-Nielsen and Shephard (2001) and Graham et al. (2017).

##### Definition 1

(Gamma and exponential distribution). *The gamma distribution is a continuous distribution on* [0,∞), *parameterized through a shape parameter α* > 0 *and rate parameter β* > 0 *(which is the inverse of the often-used scale parameter) and denoted as*

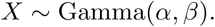

*The probability density function of X reads*

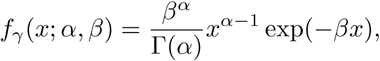

*where* 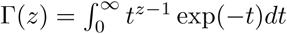 *for z* > 0 *is the gamma function. The characteristic function is given by*

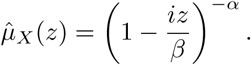

*For α* = 1, *one obtains the exponential distribution*.

##### Definition 2

(Beta distribution). *The standard beta distribution is a continuous distribution on* (0, 1), *parameterized through a shape parameter α* > 0 *and scale parameter β* > 0. *The state space can be generalized from* (0, 1) *to* (*a, c*) *by introducing the minimum and maximum values a and c as additional parameters. The resulting four-parameter distribution is denoted by*

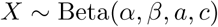

*and has probability density function*

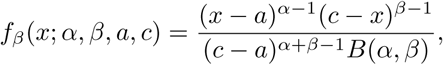

*where 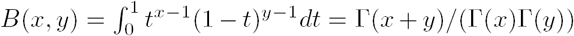 for x, y* > 0 *is the beta function. The characteristic function of the beta distribution is given by*

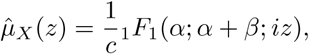

*where* _1_*F*_1_ *is the confluent hypergeometric function of the first kind (see Definition 6)*.

##### Definition 3

(Negative binomial distribution, NB). *The negative binomial (NB) distribution is a discrete distribution that describes the probability of an observed number of failures*

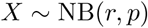

*in a sequence of independent Bernoulli trials until a predefined number of successes has occurred. In each trial, the probability of success is denoted by p* ∈ [0, 1], *and the predefined number of successes is r* ∈ ℕ_0_, *respectively. The probability mass function of X is given by*

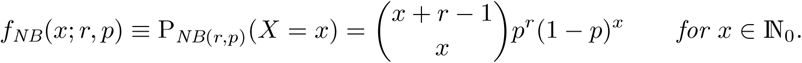

*The probability generating function of X is given by*

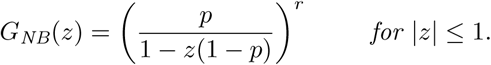

*The above definition of the negative binomial distribution can be extended to r* ∈ ℝ_+_. *All equations remain valid except for the interpretation in terms of Bernoulli trials. This generalization of r is underpinned by the construction of the Poisson-gamma distribution that is of central interest in this work and derived along Definition 5*.

*Note: Here, we describe X to represent the number of failures. Literature also provides different parameterizations, where X e. g. denotes the total number of trials (including the last success). The notation used here is the one implemented in the R function nbinom (package stats), with r and p being called size and prob. Another commonly specified parameter is the mean mu of X, given by mu* = *size/prob - size*.

##### Definition 4

(Geometric distribution). *The geometric distribution is a discrete distribution that describes the probability of*

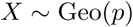

*failures before the first success in independent Bernoulli trials with success probability p each. The probability mass function of X is given by*

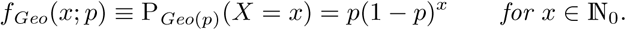

*Note: f*_*NB*(*r,p*)_(*x*; 1, *p*) ≡ *f*_*Geo*(*p*)_(*x*; 1 − *p*).

##### Definition 5

(Poisson distribution and conditional Poisson distribution). *The Poisson distribution is a discrete count distribution, denoted by*

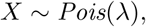

*with probability measure*

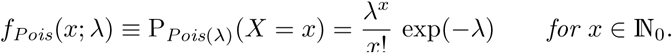

*The probability generating function of X reads*

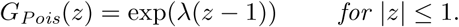

*A conditional Poisson distribution is a Poisson distribution with intensity parameter λ following itself a distribution with probability density function g, parameterized by θ. We denote this by*

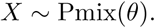

*The probability mass function of X is given by*

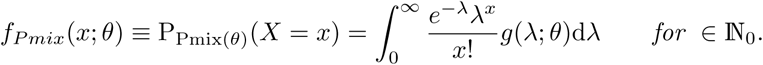

##### Definition 6

(Confluent hypergeometric function of first order). *Let w, z, a, b* ∈ℂ *Kummer’s equation*

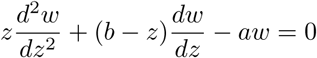

*has a regular singularity at the originand an irregular singularity at infinity. One standard solution of this differential equation that only exists if b is not a non-positive integer is given by the Kummer confluent hypergeometric function M* (*a, b, z*) *with*

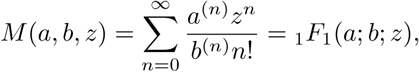

*where* _1_*F*_1_ *is the confluent hypergeometric function of the first kind with the rising factorial defined through*

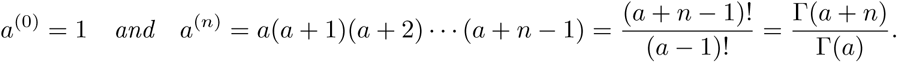

*The generalized hypergeometric function is given by*

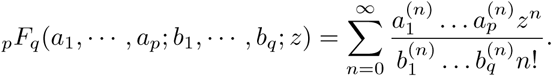

*If Re*(*b*) > *Re*(*a*) > 0, *M* (*a, b, z*) *can be represented as an integral*

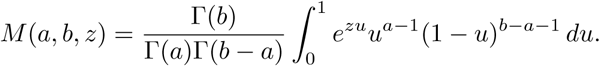

##### Definition 7

(Lévy process, subordinator). *A process* (*X*_*t*_)_*t*≥0_ *with values in* ℝ ^*d*^ *is called a Lévy process (or process with stationary independent increments) if it has the following properties:*

- *For almost all ω in the considered probability space, the mapping t* ↦ *X*_*t*_(*ω*) *is right-continuous on* [0, ∞],
- *for* 0 ≤ *t*_0_ < *t*_1_ < …< *t*_*n*_, *the random variables Y*_*j*_ := *X*_*t_j_*_ − *X*_*t_*j*-1_*_ (*j* = 1, …, *n*) *are independent*,
- *the law of X*_*t*+*h*_ − *X*_*t*_ *depends on h* > 0, *but not on t*.

*An increasing Lévy process is called a* subordinator. *Examples for Lévy processes are Brownian motion or a compound Poisson process (see Definition 8)*.

##### Definition 8

(Poisson process and compound Poisson process, CPP). *A Poisson process X*_*t*_ *with intensity parameter λ starts almost surely in zero, has independent increments, and for all* 0 ≤ *s* < *t one has X*_*t*_ *− X*_*s*_ ∼ *Pois*((*t* − *s*)*λ*). *A compound Poisson process Z*_*t*_ *with intensity parameter λ is defined as*

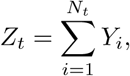

*where N*_*t*_ *is a Poisson process with parameter λ, and Y*_*i*_ *are independent and identically distributed random variables. The characteristic function of a CPP depends on the distribution of the Y*_*i*_ *and is given by*

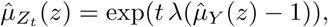

*where 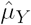 is the characteristic function of the Y*_*i*_.

##### Definition 9

(Ornstein-Uhlenbeck (OU) process). *Following Barndorff-Nielsen and Shephard (2001), an Ornstein-Uhlenbeck (OU) process y*_*t*_ *is the solution of a stochastic differential equation (SDE) of the form*

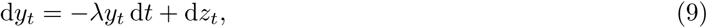

*where z*_*t*_, *with z*_0_ = 0 *almost surely, is a Lévy process (see Definition 7). If the Lévy process has no Gaussian components, the process z*_*t*_ *is called a non-Gaussian OU process or also a process of OU-type. Often, this is shortened to OU process. Barndorff-Nielsen* et al. *(2001) also call z*_*t*_ *a background-driving Lévy process (BDLP) as it drives the OU process. A special property of OU processes is that, given a one-dimensional distribution* 𝒟, *there exists an OU–type stationary process whose one-dimensional law is* 𝒟 *if and only if* 𝒟 *is self-decomposable*.

*In most applications in financial mathematics, the SDE* (9) *is transformed to*

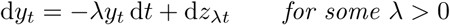

*such that whatever value of λ is chosen, the marginal distribution of y*_*t*_ *remains unchanged. In the context of our work, we however need to work with the original, untransformed SDE (9). In that case, the procedure to find* 𝒟 *for a given Lévy subordinator z*_*t*_ *is given as follows (as also described in the main text with model-specific notation):*

*1. Find the characteristic function 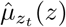 of the Lévy subordinator z*_*t*_.

*2. Calculate 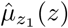 and write the result in the form* exp(*ϕ*(*z*)) *for some function ϕ*(*z*).

*3. Calculate the characteristic function C*(*z*) *of the stationary distribution* 𝒟 *of y*_*t*_ *by setting 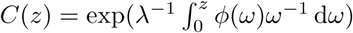. C*(*z*) *leads to* 𝒟.

*An example is shown later for the derivation of the steady-state distribution of the basic model (Figure 1A)*.

##### Definition 10

(Self-decomposable distributions). *Let 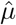 be the characteristic function of a random variable X following the one-dimensional law* 𝒟. 𝒟 *is self-decomposable iff*

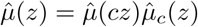

*for all z* ∈ℝ *and all c* ∈ (0, 1) *and some family of characteristic functions 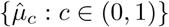*.

**Lemma** The following identities will be used in the derivations on the following pages:

1. For the gamma function Γ, one has

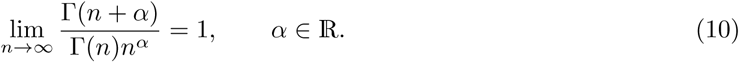
2. Using

- the identity of the binomial series theorem:

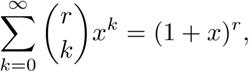

which holds for |*x*| < 1 and *r* can be arbitrary real or complex,
- the symmetry of binomial coefficients

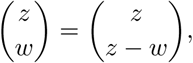

with *z* ∈ℝ > *w* ∈ℝ *≥* 0,
- and the identity for upper negation of binomial coefficients

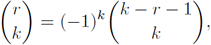

with an integer *k*,
one has

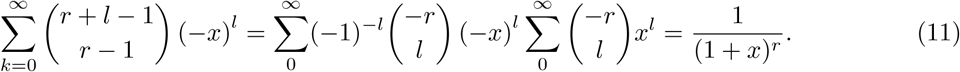

Here, *r* can be any arbitrary real or complex number but |*x*| < 1.

#### NEGATIVE BINOMIAL CORRESPONDS TO POISSON-GAMMA

Negative binomial and Poisson-gamma distributions are equivalent, i. e. they can be transformed into each other by reparameterization. To show this, we start with a Poisson-gamma (PG) distribution. Let *α, β* > 0 and *x* ∈ℕ_0_. Then, according to Definitions (1) and (5),

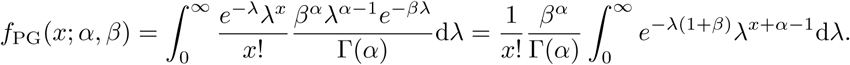

Substitution with *u* = *λ* (1 + *β*) and 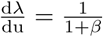 and use of 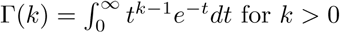 for *k* > 0 leads to

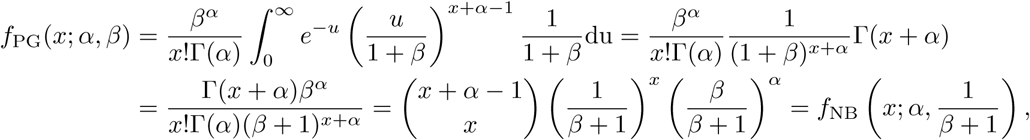

which is the probability mass function of the negative binomial distribution. The reparameterization can also be considered the other way round:

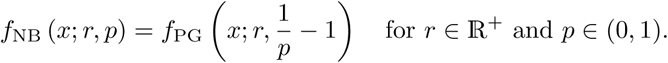

#### POISSON-BETA CONVERGES TOWARDS NEGATIVE BINOMIAL

In the Results section, we considered the Poisson-beta distribution PB (*r*_*act*_ */r*_*deg*_, *r*_*deact*_ */r*_*deg*_, 0, *r*_*on*_ */r*_*deg*_) (see Definitions 2 and 5) as the steady-state distribution of the switching model. For large *r*_*deact*_ */r*_*deg*_ and *r*_*on*_ */r*_*deact*_ < 1, the probability mass function of this distribution converges towards the one of a negative binomial distribution (see Definition 3) (Raj et al., 2006):

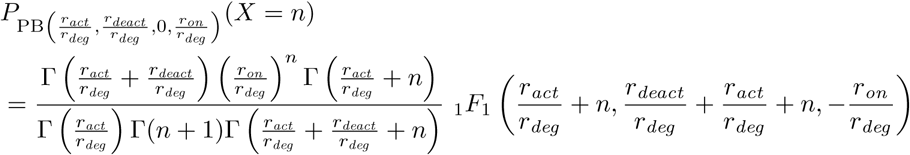

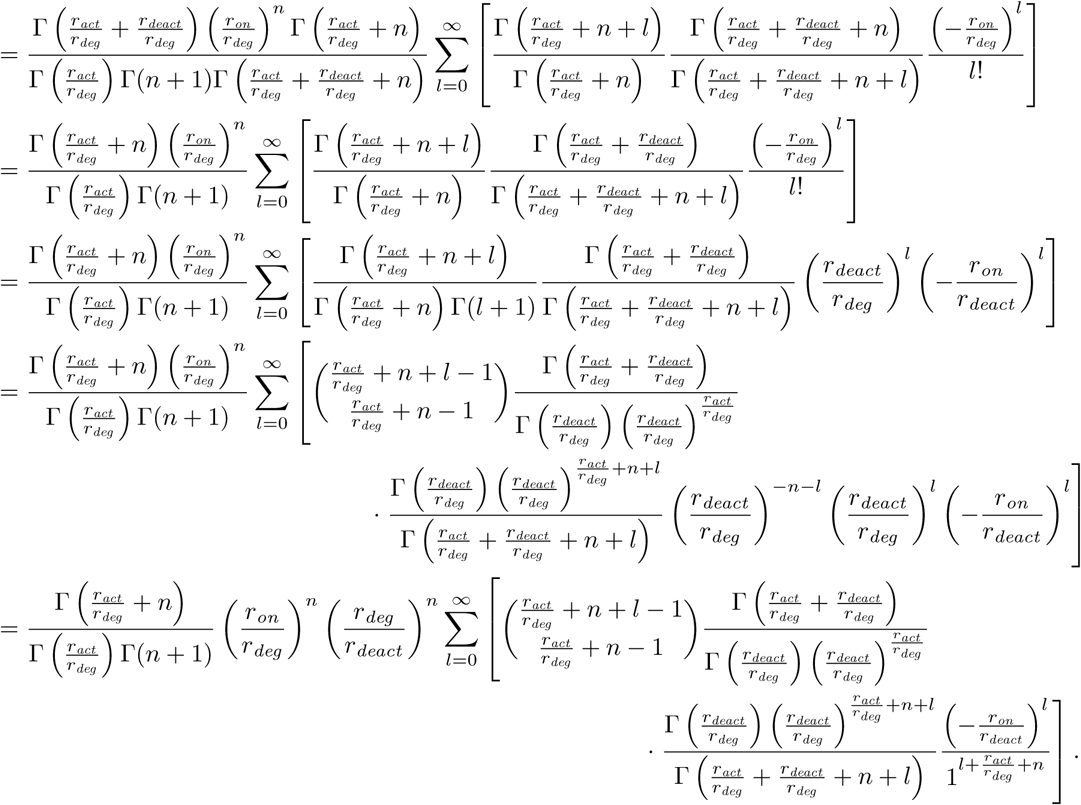

Taking the limit, one can use the asymptotic approximation given in (10) twice, leading to

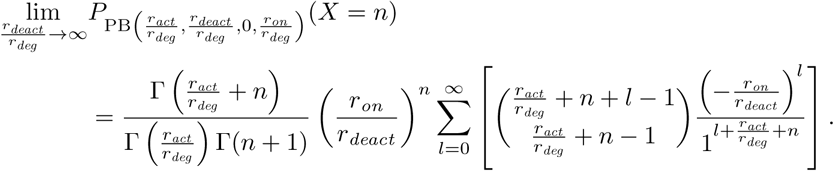

Next, we use (11) to simplify the expression, and to that end assume *r*_*on*_ */r*_*deact*_ < 1:

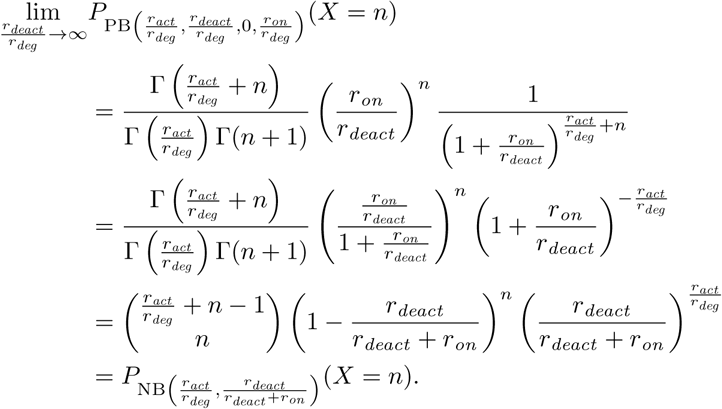

This is the probability mass function of the negative binomial distribution NB (*r*_*act*_ */r*_*deg*_, *r*_*deact*_ */r*_*deact*_ + *r*_*on*_). Overall, for *r*_*deact*_ */r*_*deg*_ → ∞ and *r*_*on*_ */r*_*deact*_ < 1, one obtains

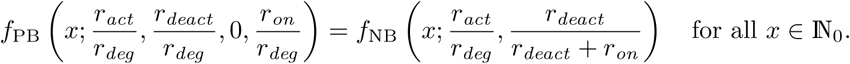

#### MASTER EQUATION OF THE GENERALIZED MODEL

We describe the derivation of steady-state distributions for mRNA counts in the considered mechanistic transcription and degradation models depicted in Figure 1, starting with the generalized model. In the following, *P* (*n, t*) describes the probability of having *n* mRNA molecules at time *t* in the system. The master equation is set up by looking at the reactions (at most one) that can happen within an infinitesimally small time interval: Either one mRNA molecule is transcribed, which happens with probability rate *R*_*t*_, or one mRNA molecule degrades with rate *r*_*deg*_, or nothing happens. In the following, we write 𝒫 (*n, t*|*R*_*t*_, *r*_*deg*_) = 𝒫 (*n, t*) for the sake of simpler notation. The master equation reads

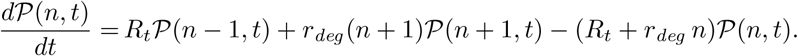

From this, one obtains the probability generating function

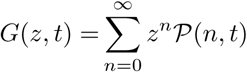

with derivatives

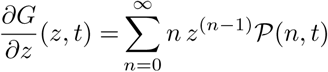

and

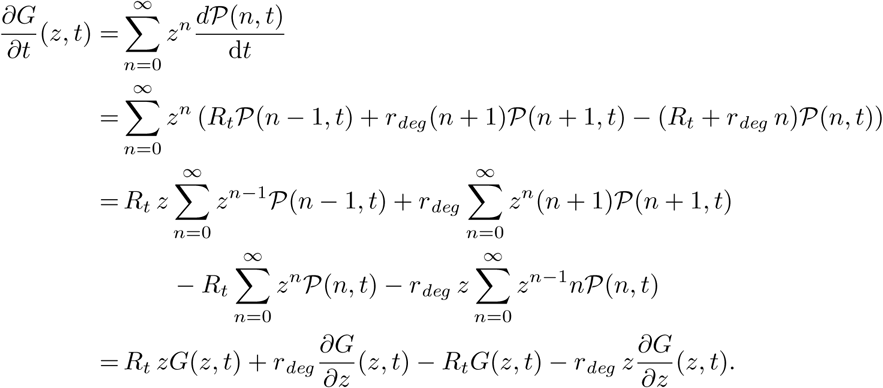

This results in the partial differential equation (PDE)

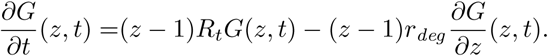

The solution of this PDE with initial condition of having *n*_0_ mRNA molecules is calculated by using the methods of characteristics:

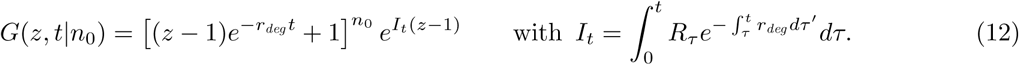

The first factor of *G*(*z, t*|*n*_0_) reflects the dependence of the distribution on the initial value *n*_0_. The second factor exp(*I*_*t*_(*z* − 1)) corresponds to the long-term behaviour of the mRNA content and equals the time-dependent probability generating function of a Poisson distribution with intensity parameter *I*_*t*_ (see Definition 5). One commonly considers the distribution in steady state (if that state exists), meaning *t* → ∞. In this limit, the first factor vanishes (i. e. becomes one). Thus, the steady-state distribution is independent of the starting condition. The second term remains. Thus, in steady state the mRNA count follows a conditional Poisson distribution with intensity parameter *I*_*t*_ being governed by the transcription and degradation process. From Definition 5, one gets

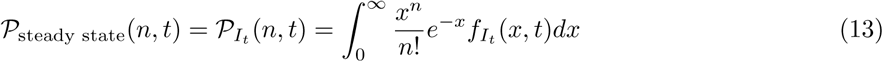

for *n* ∈ ℕ_0_ and *t* ≥ 0 (but large), where 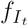 denotes the density of *I*_*t*_. To exactly specify the conditional Poisson distribution we need to take a closer look at the intensity process *I*_*t*_, defined through (12), and examine its long-term (steady-state) behavior. 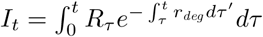 is a solution of the random differential equation (RDE)

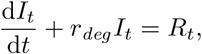

which can be rewritten as

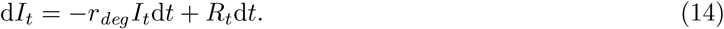

In this representation, one can directly recognize the impact of the mRNA degradation rate *r*_*deg*_ and the transcription rate *R*_*t*_ on the number of mRNA molecules: Larger *r*_*deg*_ will lead to lower mRNA numbers, larger *R*_*t*_ to higher numbers. The properties and steady state of *I*_*t*_ clearly depend on the choice of *R*_*t*_. The RDE (14) can be generalized to a stochastic differential equation by considering *R*_*t*_d*t* = d*L*_*t*_, where *L*_*t*_ is an arbitrary (increasing) Lévy process (Definition 7). Then

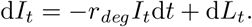

Since the trajectories of a Lévy process are not necessarily left-continuous, their derivatives may not exist in the classical sense. Care has to be taken here (see main text).

In the following sections, we show how to derive the steady-state distribution of *I*_*t*_ for different choices of *R*_*t*_ or *L*_*t*_.

#### DETERMINISTIC CONTINUOUS TRANSCRIPTION MODEL

We start with a simple model: If *R*_*t*_ is a deterministic rather than stochastic function *R*(*t*), *I*_*t*_ itself becomes deterministic, now denoted by *I*(*t*). Dattani and Barahona (2017) show that the probability to have *n* mRNA molecules at time *t* is Poisson distributed with time-dependent intensity *I*(*t*), i. e.

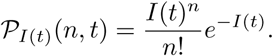

The solution for *I*(*t*) then is

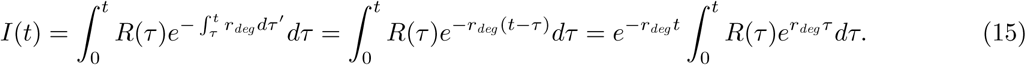

#### BASIC MODEL

In the basic model (Figure 1B), *R*(*t*) takes only one time-independent value *r*_*tran*_. With Equation (15), we get

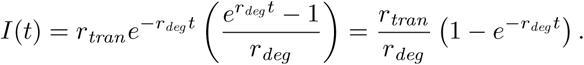

All together, for *t* → ∞, the steady-state distribution of the mRNA count follows a Poisson distribution with intensity parameter *I* = *r*_*tran*_ */r*_*deg*_.

#### SWITCHING MODEL

We now assume transcription to be governed by *R*_*t*_ = *r*_*switch*_ (*t*), which is a Markov chain with two states *on* (or *active*) and *off* (or *inactive*), switching between these two states after exponentially distributed waiting times with rates *r*_*act*_ and *r*_*deact*_. During the *active* state, transcription happens with rate *r*_*on*_, whereas in the *inactive* state, either strongly down-regulated transcription happens (small *r*_*off*_) or none (*r*_*off*_ = 0). Supplementary Figure S1 shows a more detailed picture of Figure 1C.

Again we calculate the steady-state distribution of mRNA content. We follow the derivation of Smiley and Proulx (2010), who show how to obtain the density function for the mRNA expression level. Dattani and Barahona (2017) use this result and transfer it into the probability distribution. Raj et al. (2006) arrive at the same solution.

The differential equation (14) now reads

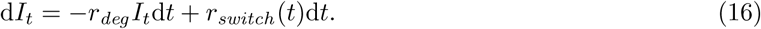

As transcription is governed by a Markov chain which is a random process and not deterministic anymore, the probability distribution for the amount of mRNA at time *t* is a Poisson mixture distribution as described by (13). Again, in order to determine the steady-state distribution of mRNA counts, we need to determine the steady-state distribution of *I*_*t*_ in (16).

The Markov chain *r*_*switch*_ (*t*) can be characterized by its infinitesimal generator

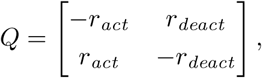

where the entries on the anti-diagonal *Q*_*ij*_ (*i* ≠ *j*) are the transition rate constants from state *j* to *i* and its reciprocals are the means of the exponential waiting times. States 1 and 2 correspond to the inactive and the active state, respectively. This means *r*_*act*_ corresponds to the rate with which a gene is activated (transition from state 1 to 2), and *r*_*deact*_ is the deactivation rate, that is the rate of the transition from state 2 to 1. The probability transition matrix *P* (*t*) is defined as

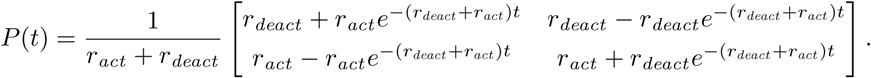

*P* (*t*) satisfies the Kolmogorov differential equation *P* ′ (*t*) = *QP* (*t*), and the initial condition is

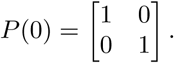

The entry *P*_*ij*_(*t*) denotes the probability of a transition from state *j* to *i*. (Note: Here, *Q* and *P* (*t*) are the transpose of the usual notation as this notation is more convenient in the present stationary analysis.) If the probabilities for *r*_*switch*_(0) being in state 1 or 2 are given by *p*(0) = [*p*_off_(0), *p*_on_(0)]^*T*^, then the distribution of *r*_*switch*_ (*t*) is given by *p*(*t*) = *P* (*t*)*p*(0) and it follows that

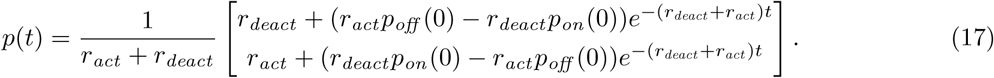

The matrix *p*(*t*) has to fulfill the Kolmogorov differential equation

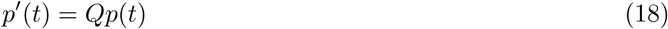

as well. Assume 0 ≤ *r*_*off*_ < *r*_*on*_, then *I*_0_ ∈ [*r*_*off*_ */r*_*deg*_, *r*_*on*_ */r*_*deg*_] and, with probability one, one has *I*_*t*_ ∈ [*r*_*off*_ */r*_*deg*_, *r*_*on*_ */r*_*deg*_] for *t* > 0. One has

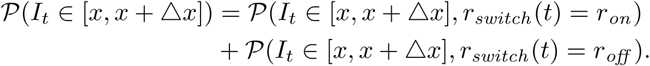

The joint cumulative distribution functions (CDFs) associated with the joint probabilities of *I*_*t*_ being equal to *x* and *r*_*switch*_ (*t*) being equal to *r*_*i*_ are given by

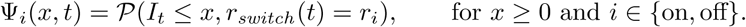

Their derivatives with respect to *x* given the joint distribution of *I*_*t*_ = *x* and *r*_*switch*_ (*t*) = *r*_*i*_ is denoted as *ψ*_*i*_(*x, t*). The probability density function (PDF) *ψ*(*x, t*) associated with *I*_*t*_ can be characterized by a system of two partial differential equations (PDEs)

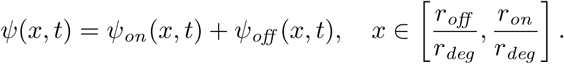

Clearly, with (17) one obtains

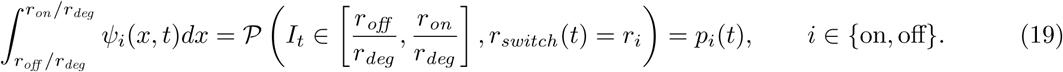

We now set

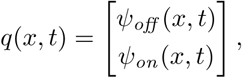

which is still directly connected with the two-state Markov chain *r*_*switch*_ (*t*). Both components of *q*(*x, t*) are continuous PDFs, one for each state of *r*_*switch*_ (*t*). This is again a two-state Markov chain and adopts the transition rate matrix *Q* from the process *r*_*switch*_ (*t*). It hence inherits its property (18), and thus, *q*(*x, t*) fulfills the Kolmogorov differential equation as well, i. e.

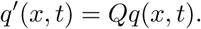

All together

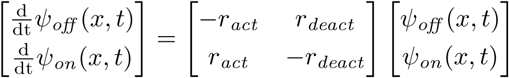

and thus

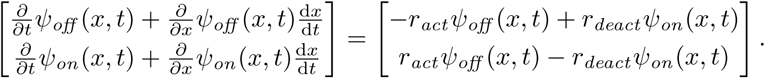

Using (16), we get 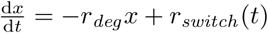. Plugging this in, the system of PDEs can be simplified to

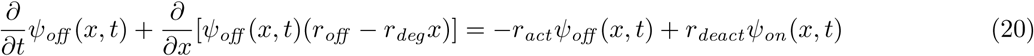

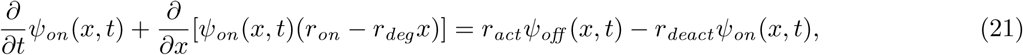

which correspond to Equations (6) in Smiley and Proulx (2010). Integrating both sides of (20) and (21) with respect to *x* over the range from *r*_*off*_ */r*_*deg*_ to *r*_*on*_ */r*_*deg*_ leads us to

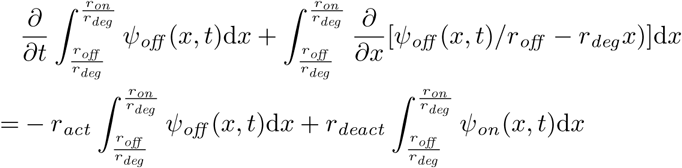

and

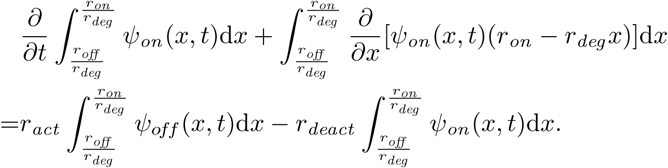

With (19), it follows that

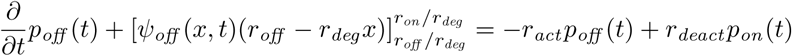

and

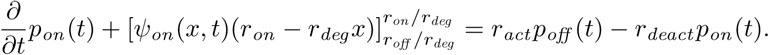

Since Equation (18) still has to be fulfilled, it follows directly that the redundant terms have to be equal to zero:

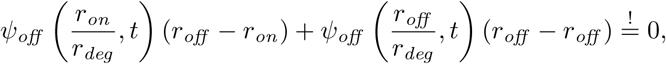

which is equivalent to

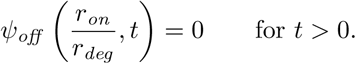

Similarly,

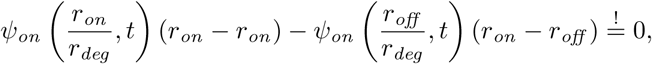

which implies

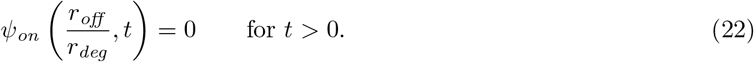

Following Smiley and Proulx (2010), we next investigate the PDF of the stationary distribution of *ψ*(*x, t*), denoted by 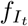, which is analogously determined by a pair of functions 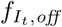 and 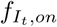 via

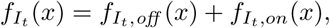

with 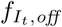 and 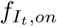 being the time-independent solutions of (20) and (21). Those can be calculated by solving the time-independent versions of (20) and (21), given by

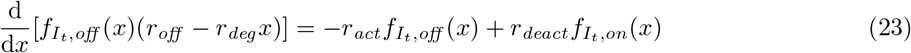

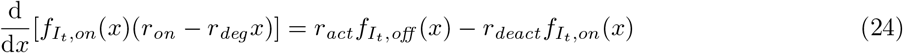

with integral conditions derived from Equation (19) for *t → ∞*

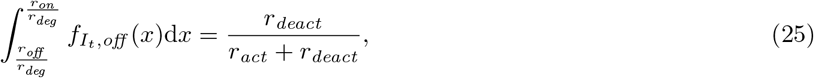

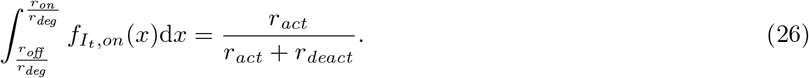

Summing up (23) and (24) results in

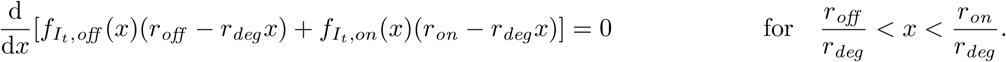

For any solution of (23) and (24) and for any constant *K* it follows that

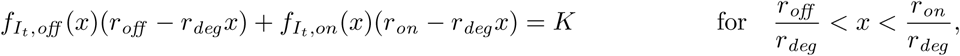

thus

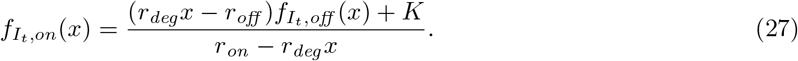

Plugging in (27) into (23) and setting *K* = 0 (as all steady-state solutions have to satisfy the condition given in (22)), we get

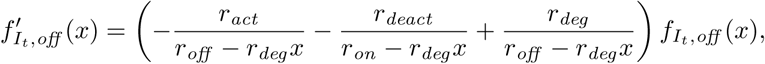

which can be solved up to a normalizing factor *C*:

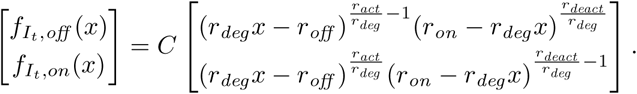

We use Equations (25) and (26) to determine *C*:

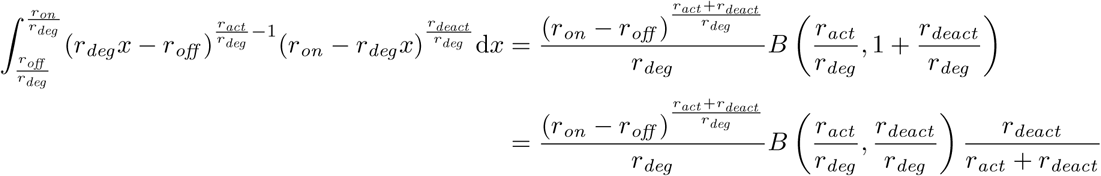

and

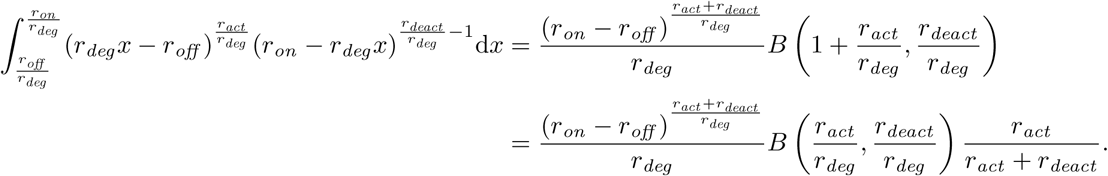

Here, *B* denotes the beta function as introduced in Definition 2. Both of the above integrals have to be normalized by

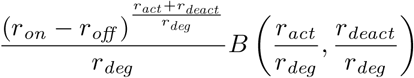

in order to result in *r*_*deact*_ */*(*r*_*act*_ +*r*_*deact*_) as given by (25) and *r*_*act*_ */*(*r*_*act*_ +*r*_*deact*_) as given by (26), respectively. All together, we get

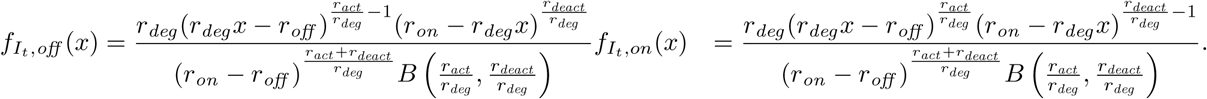

Adding these up will provide the final solution

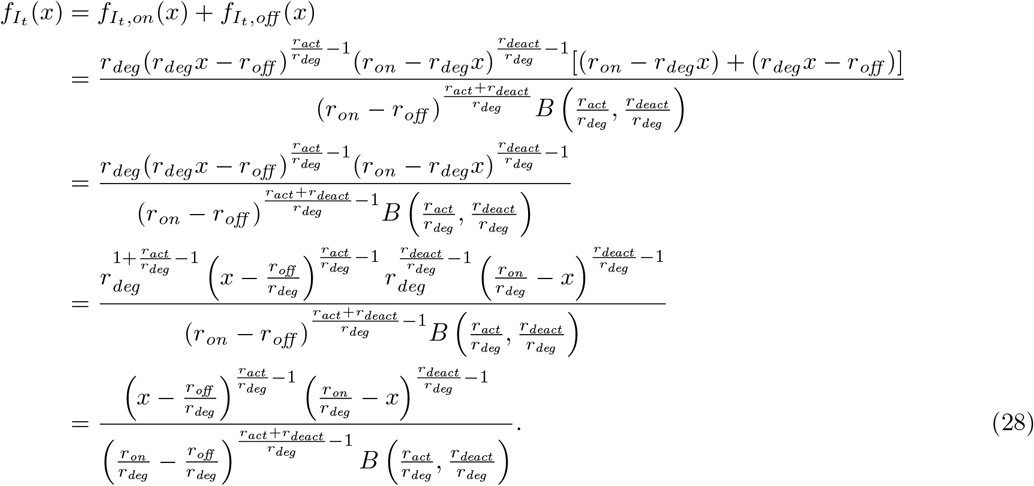

This is the density of the stationary distribution of *I*_*t*_ from Equation (3), and it is the density function of a four-parametric beta distribution (see Definition 2) with parameters *a* = *r*_*off*_ */r*_*deg*_, *c* = *r*_*on*_ */r*_*deg*_, *α* = *r*_*act*_ */r*_*deg*_ and *β* = *r*_*deact*_ */r*_*deg*_.

The overall steady-state distribution of mRNA counts (see Equation (13)) is by construction a conditional Poisson distribution. When conditioning the Poisson distribution on an intensity parameter following the distribution defined by Equation (28), the overall distribution will be a Poisson-beta distribution whose probability mass function can be written in the following way:

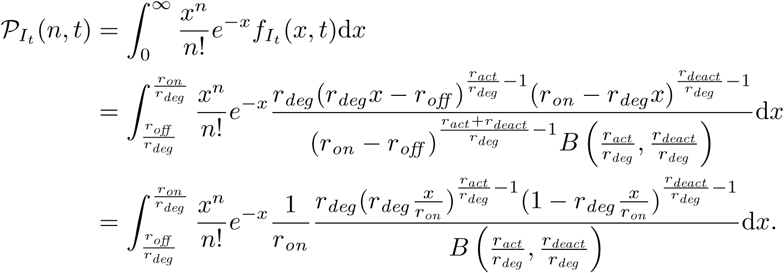

Substitution by *z* = (*xr*_*deg*_ − *r*_*off*_*)/*(*r*_*on*_ − *r*_*off*_) and d*x/*d*z* = (*r*_*on*_ − *r*_*off*_*)/r*_*deg*_ leads to

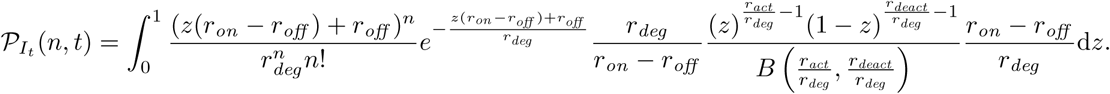

From this point on, we can simplify further when setting *r*_*off*_ = 0. This is valid as we suppose no transcription during the deactivated DNA state. Then

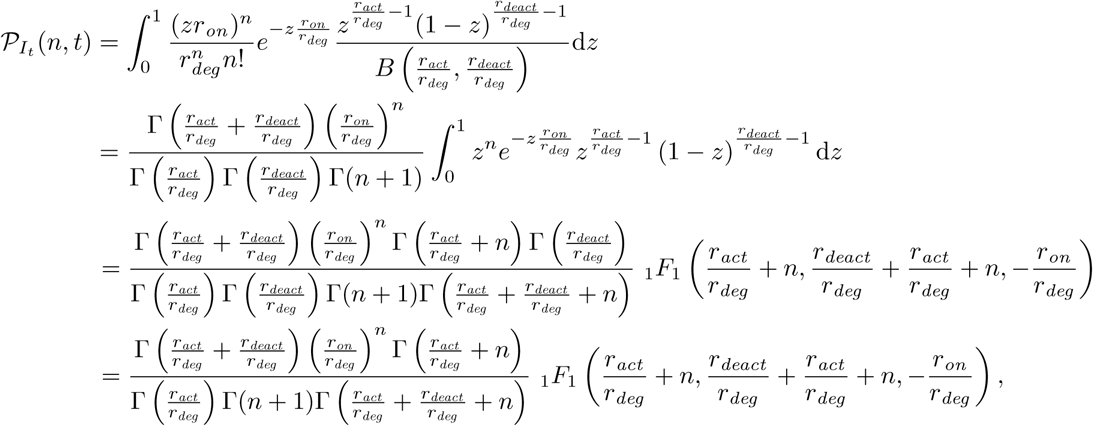

where _1_*F*_1_ is the confluent hypergeometric function of first order as introduced in Definition 6.

#### OU PROCESSES LINK SDES TO STEADY-STATE DISTRIBUTIONS

OU processes and the concept of linking them to distributions is widely used in financial mathematics, especially in the areas of option pricing and volatility modeling. Among others (Sato (1999), Rogers and Williams (2000)), especially Barndorff-Nielsen and Shephard (2001) and Barndorff-Nielsen et al. (2001) used OU processes in a wide range and showed and proved a substantial amount of their properties.

#### OU PROCESS DERIVATION FOR BASIC MODEL

In the following, we will show how to use an OU process to infer the steady-state distribution of the basic model (Figure 1A). We use the following general OU equation introduced in the main text in Equation (5):

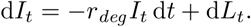

This general OU SDE is transformed to the ODE of the basic model by setting *L*_*t*_ := *r*_*tran*_ *t*, with d*L*_*t*_ = *r*_*tran*_ d*t*, yielding the ODE

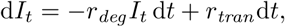

which was already given in the main text as Equation (2). In this simple case, the Lévy subordinator *L*_*t*_ = *r*_*tran*_*t* describes a state-continuous process without any jumps or Brownian components. Still, this ODE fulfills all required properties and can be used for deriving a steady-state distribution for the mechanistic model according to the procedure that was described before.

To do so, we now follow the three steps described in Definition 9 and in the main text. These are:

1. Find the characteristic function of the Lévy subordinator *L*_*t*_ = *r*_*tran*_ *t*. For the basic model, that is

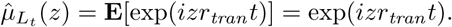
2. Calculate 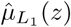 and write the result in the form exp(*ϕ*(*z*)) to determine *ϕ*(*z*). For the basic model, that is

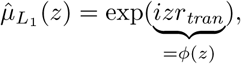

so it follows that *ϕ*(*z*) = *izr*_*tran*_.
3. Calculate the characteristic function *C*(*z*) of the stationary distribution *gD* of *I*_*t*_ by

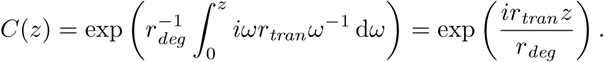

This is the characteristic distribution of a point distribution where all mass is concentrated at a single point *r*_*tran*_ */r*_*deg*_ (see Sato (1999), Example 2.19). This is also the same solution that we obtained by solving the ODE directly, shown in the previous sections.

#### MASTER EQUATION OF THE BURSTING MODEL

When the mechanistic model of the bursting process in known, its master equation can be set up easily, especially if one draws a connection to queuing theory. In a general queuing model, customers arrive at one or several service desks according to some arrival process, which in our case corresponds to the transcription process. The number of customers waiting is equivalent to the number of mRNA molecules in a cell. As soon as a customer can proceed from the queue to a service desk, this number decreases by one, corresponding to mRNA degradation. Here, service time is zero and thus plays no role in our model.

The bursting model described in the main text corresponds to the following queuing system: Customers do not arrive separately at constant rate, but they arrive in groups (e. g., in buses) after exponentially distributed waiting times with rate *r*_*burst*_. Then, several people start queuing at the same time. The number of people arriving with each group follows a geometric distribution with mean *s*_*burst*_.

This process corresponds to a mixture of two queuing problems from Adan and Resing (2002). The first queuing problem is the basic so-called *M/M/*∞ queuing setup (Example 11.1.1 in that reference), and the second one is the *M/G/*1 model which corresponds to a queue with group arrivals (Chapter 10.4 in that reference). (The notation here is due to Kendall: In the three-part code *a/b/c, a* specifies the inter-arrival time distribution, *b* the service time distribution and *c* the number of servers. The letter *G* is used for a general distribution, *M* for the exponential distribution and *D* for deterministic times.) A standard waiting process is modeled where the group arrival time is exponentially distributed, service time and group size follow arbitrary distributions, but only one service counter is open. With those two models in mind, we set up our bursting queuing process (as mentioned above we don’t have service times). We illustrate all possible state changes in Supplementary Figure S2. Along that figure, we can set up the master equation directly:

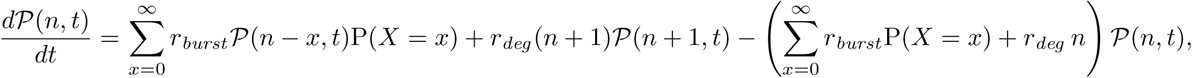

where P denotes the probability mass function of a random variable *X* that is geometrically distributed with success probability *p*. The probability-generating function then reads

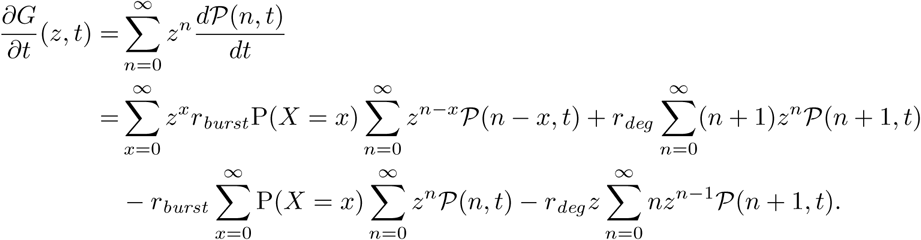

With

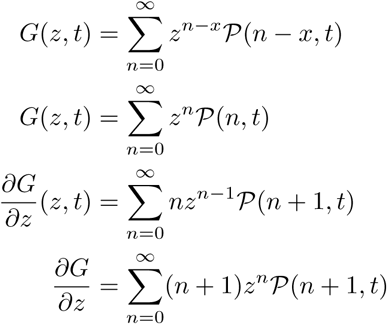

it follows that

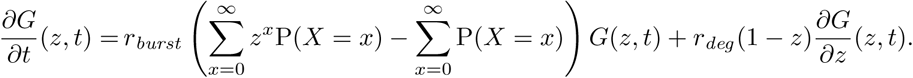

Because of P(*X* = *x*) = (1 *- p*)^*x*^*p* and 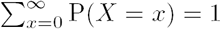, it follows that

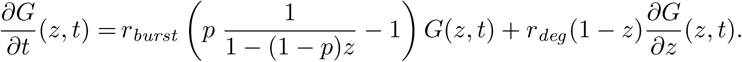

Taken together, the result is a PDE of order one and equivalent to

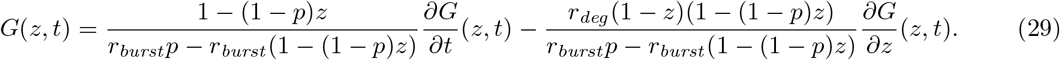

In the following we show how to solve the PDE

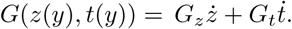

*Ansatz: 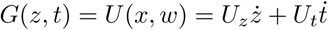*

To use this ansatz, we need to determine *x* and *w*. We read *ż* and 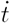 from the full equation given by (29):

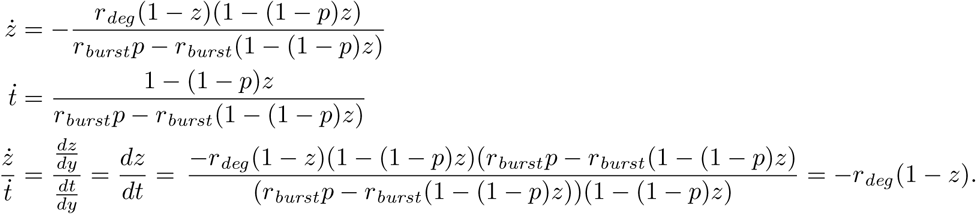

Thus, it follows that

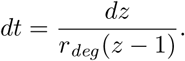

Integrating both sides yields

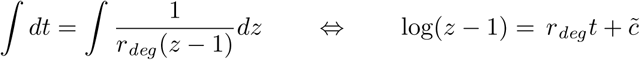

for an arbitrary constant 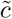. Next, we take the exponential of both sides

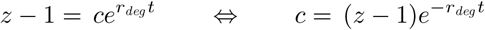

for a constant *c*. Choose 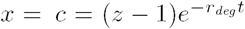 and *w* = *t*. Then it follows that 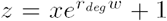. For the derivatives, we obtain

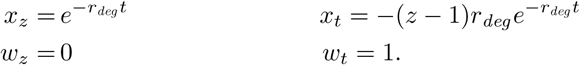

Next, we need to determine *U*_*z*_ and *U*_*t*_:

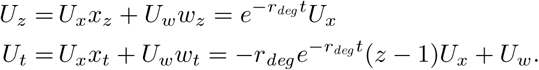

Finally, we can compute U:

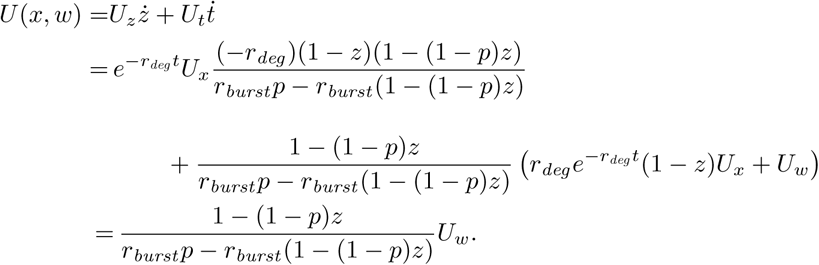

Plug in *z* and *t* to get *U* only in terms of *x* and *w*:

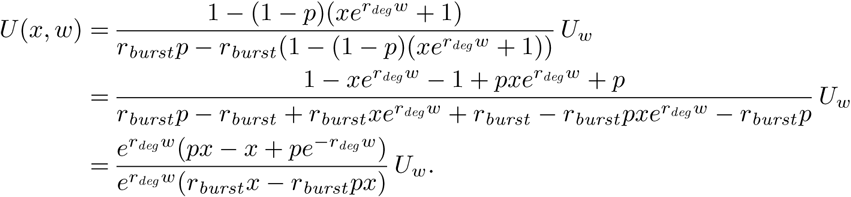

As *U*_*w*_ = *dU/dw*, we can separate the terms depending on *U* and the terms depending on *w*:

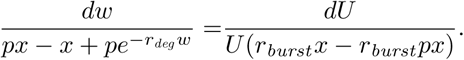

Integrating both sides leads to:

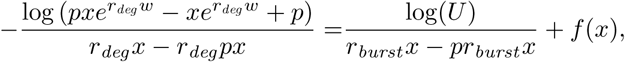

where *f* (*x*) is seen as a constant with respect to *w* and *U* and thus can only be a function that depends on *x*. Then

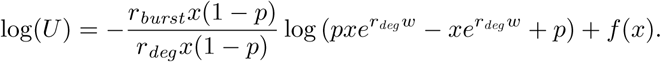

Next, we take the exponential on both sides

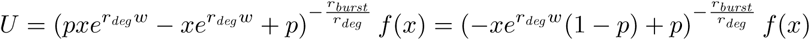

Return to the parameterization in terms of *z* and *t*:

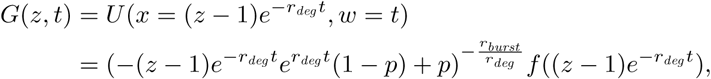

where 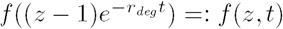 now represents a function that depends on *z* and *t*. We get

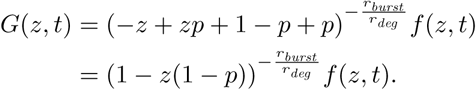

The right hand side is of the form of the probability generating function of a negative binomial distribution with parameters *r*_NB_ and *p*_NB_ as stated in Definition 3 if one chooses 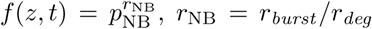 and *p*_NB_ = *p*. Since the mean burst size in the bursting model is *s*_*burst*_, the parameter *p* of the geometric distribution and hence the parameter *p*_NB_ of the negative binomial distribution is equal to (1 + *s*_*burst*_)^*-*1^.

#### R PACKAGE scModels

We need to calculate the probability mass function of the Poisson-beta distribution (Equation (4)) in some sections of this paper. The general form of the probability mass function of the Poisson-beta(*α, β*, 0, *c*) distribution for *α, β, c* > 0 is given by

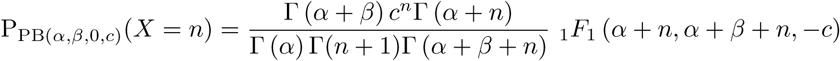

for *n* ∈ IN_0_. To compute this function, the Kummer function _1_*F*_1_ (*a, b, z*) (see Definition 6) needs to be calculated with the following constraints on its parameters:

1. *z* ∈ ℝ_≤0_ (where *z* is the third parameter of _1_*F*_1_); in our case where *z* = −*c*, it thus follows 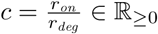
2. *a, b* ∈ ℝ_≥0_ and 0 ≤ *a* ≤ *b*. This means in our case where *a* = *α* + *n* and *b* = *β* + *n* that 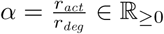 and 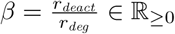 and *n* ∈ ℤ_≥0_.

Muller (2001) showed how hard it is to compute the Kummer function, because its computational behaviour splits into a number of distinct regions, which makes it impossible to have a unified algorithm for all possible input parameters. One of the well-behaved analytical solutions to the function is in the form of an infinite series. Additionally, for specific constraints on the parameters (which are fullfilled when the function appears inside the Poisson-beta distribution), there exists an integral representation of the solution. Nevertheless, neither the integral nor the infinite sum can be computed directly, and thus approximations and workarounds had to be implemented. There are different existing methods that have tried to address this problem. On the one hand, there are methods that compute the Poisson-beta distribution by approximating the integral representation of the Kummer function (see **BPSC**, Vu et al., 2016); while on the other hand methods employ the characteristics of the Poisson-beta distribution to estimate its parameters, circumventing the evaluation of the Kummer function (see **D3E**, Delmans and Hemberg, 2016). Our approach is to calculate the density by truncating the infinite series solution to the Kummer function at a reasonable error bound. This is also challenging as the existing R function kummerM() (Package:**fAsianOptions**) tries a similar approach but fails for many parameters (see Supplementary Figure S3). In the following, we will first go into detail of the existing methods and will then present our new implementation. Afterwards we compare our method to existing ones in terms of fitting and computation time.

#### BPSC

Vu et al. (2016) present how to do use the integral representation to calculate the probability mass function of a Poisson-beta distribution. This is implemented in their R-package **BPSC**. Vu et al. (2016) define three different beta-Poisson models (they use this name rather than Poisson-beta) where the so-called three-parameter beta-Poisson model corresponds to the one we proposed in the main part of this paper, and thus, is the only one we want to use here and later on in the comparison. Parameter estimation is done via likelihood maximization, where two techniques are used to speed up the calculations: First, the authors bin the data and for each bin interval the probability is calculated separately via the PDF of the Poisson-beta distribution in this interval. Second, to calculate the PDF of such an interval, the integral-notation of the Kummer function is used and the value of this integral is approximated by using the Gaussian quadrature method. Starting values for *α* and *β* for the parameter optimization are calculated based on the method of moments whereas *c* is assumed to be the maximum of the data points.

#### D3E

Delmans and Hemberg (2016) implemented two different methods to estimate the parameters of the Poisson-beta distribution in their **D3E** package that is available in Python: The first implementation is a “fast but inaccurate method” using the moment matching approach that was first proposed by Peccoud and Ycart (1995). The second implementation is the Bayesian inference method proposed by Kim and Marioni (2013) where gamma priors are used for the parameters *α, β* and *c* and a collapsed Gibbs sampler, using slice sampling, is used for parameter estimation. Additionally, **D3E** provides a differential gene expression test by using a likelihood ratio test. To overcome the problem of calculating the Kummer function, a Monte Carlo method is used that approximates the PDF as average of empirical PDFs of a large number of datasets generated from a Poisson-beta distribution.

#### scModels

All functions needed to simulate data or estimate distributions are collected in our R package **scModels** which is published on CRAN (https://cran.r-project.org/). The current working version can be found on Github under https://github.com/fuchslab/scModels. Included are the Poisson, the negative binomial, and most importantly, a new implementation of the Poisson-beta distribution (probability density function, cumulative distribution function, quantile function and random number generation) together with a required new implementation of the Kummer function (also called confluent hypergeometric function of the first kind). Three implemented Gillespie algorithms allow synthetic data simulation via the basic, switching and bursting mRNA generating process, respectively. Lastly, we added likelihood functions for one population and two population mixtures – with and without zero inflation – that allow estimation of the Poisson, negative binomial and the Poisson-beta distribution. These can be performed with one included wrapper function fit params() that uses the general-purpose optimization function optim().

As stated above, we implemented a new version in R of the Kummer function that uses the infinite sum representation. The only existing (at least to our knowledge) implementation in R, kummerM(), which is contained in the package **fAsianOptions**, works only for some specific parameter choices but not for others, e.g. for negative *z* the kummerM() does not return the correct values (see Supplementary Figure S3). More specifically, this implementation gives back the correct result only for parameter values that can be written as 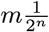 with *m, n* ∈ ℕ_0_. Because this is impracticable when numerically determining parameters during likelihood optimization, we decided to correct this issue by reimplementing the Kummer function.

Our new implementation aims to be as close as possible to the true solution for the parameter values we need, when the Kummer function is used during calculation of the Poisson-beta probability mass function. Muller (2001) stated that if neither *a* nor *b* are negative integers, then the series converges for all finite *z*. In reality, however, calculations fails when, for example, *a* and *z* have opposite signs. The problem arises because of cancellations. One of Kummer’s transformations promises to circumvent this problem: Suppose that 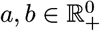 and 0 ≤ *a* ≤ *b* but *z* ∈ ℝ_*-*_, then

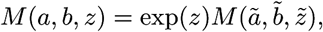

where 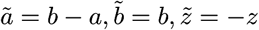 Now for the new parameters it holds that

1. 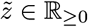.
2. 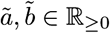 for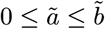.

With these new constraints, the power series does not have convergence issues, but is difficult to be evaluated because of limits on machine precision. Consequently, we use the MPFR library (see https://www.mpfr.org) for arbitrary-precision floating-point computation. To make the code more readable, we use another MPFR C++ wrapper (http://www.holoborodko.com/pavel/mpfr/), written by Pavel Holoborodko. The precision of the temporary results in an expression is chosen as the maximum precision of its arguments, and the final result is rounded to the precision of the target variable.

Although the final result of the function is quite large, the logarithmic value can be casted into double, which is then used further. We implement the iterative algorithm described as Method 1 in Muller (2001). Convergence and error analysis for Taylor series summation using multiple precision arithmetic has been explained in Brent (2010).

Convergence of the Kummer series as given in Definition 6 can be checked using the ratio test, and an appropriate lower bound on the number of terms needed for computation can be subsequently calculated. One has

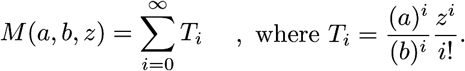

For convergence, we need

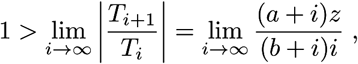

which is easily fulfilled for all reasonable positive values of *a, b, z*. With this, we can have a lower bound on the number of terms needed for a good approximation. Specifically, we need to sum up at least until the term where the ratio falls below one. Hence, the condition is

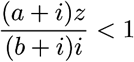

and this implies

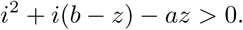

Since only positive values of *i* are sensible, we have

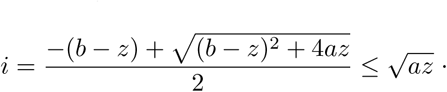

Therefore, the series converges after 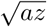 terms. Nevertheless, our new implementation of the Kummer function that is contained in **scModels** stops the calculations of the infinite sum as soon as a new summand is smaller than 10^*-*^6.

#### *COMPARISON OF* **scModels** *WITH* **D3E** *AND* **BPSC**

In a simulation study, we compare the implemented functions of the Poisson-beta distribution that are contained in the three described packages. We first generate sample data on which to test the three packages by using our gmRNA switch() function contained in **scModels**. We use this function to generate data from the switching model as this is the mechanistic model that leads to the Poisson-beta distribution in steady state. We simulate 1,000 data points from four different sets of parameters, respectively. Supplementary Table S2 shows the chosen Poisson-beta parameters which are calculated from the parameters used in the data simulation, *α* = *r*_*act*_ */r*_*deg*_, *β* = *r*_*deact*_ */r*_*deg*_ and *c* = *r*_*on*_ */r*_*deg*_, as well as the results of this comparison study. These results are also depicted in Supplementary Figure S4. The estimation procedures and time measurements were performed on a cluster of machines with the following specifications: Intel(R) Xeon(R) CPU E5620 (2.40GHz). Jobs were submitted using the Univa Grid Engine queuing system with 1 GB of memory for each job. Package-specific details of the procedure are described in the following:

- **BPSC:** The function getInitParam() estimates initial parameters of the distribution to be passed to the optimization function. The estimateBP() function calls the standard optim() routine to generate final results.
- **D3E:** D3E is designed for identifying differentially expressed genes based on scRNA-seq data. The data needs to be provided in a tab-separated read-count table, where rows correspond to genes, and columns correspond to cell types. Since it works for differentially expressed genes, the columns in the read-count table have to be labeled for the two different types of cells or conditions. The output is the parameter values of the Poisson-beta distribution along with other statistics for comparison. Here, we do not aim to test for differential expression but only intend to estimate model parameters for one type of cells. Hence, we have to circumvent this procedure: We use the function getParamsBayesian() from inside the package to bypass the differential expression step.
- **scModels:** We use the method of moments combined with bootstrap to predict initial values for the optimization. The final result is obtained by minimizing the negative log-likelihood function that employs the implemented density function dpb() of the Poisson-beta distribution.

The estimation results show that all three methods are able to estimate a density function that well describes the data and is close to the true density curve (Supplementary Figure S4). The obtained values of the negative log-likelihood are in the same range, with our package **scModels** always leading to the lowest or equally low (i.,e., best) value (Supplementary Table S2). Computing times and parameter estimates are variable and do not show a clear picture.

#### DATA APPLICATION

In the main text, investigations were performed on publicly available real-world datasets. Here we describe some of the (additional) analysis in more detail.

#### GENE FILTERING

The data preprocessing has been performed as follows.

##### Nestorowa dataset

(Nestorowa et al., 2016). As described in the main text, this data was generated using the Smart-Seq2 protocol and thus the resulting data consists of read counts. The original data matrix contained 45,771 genes and 1,656 cells. We used two filters: The first one selects only those genes that have mean expression larger than one, whereas the second filter additionally removes all genes that are only lowly expressed, i.e. after application of this filter, only those genes remain that have at minimum five reads in at minimum 20 cells. After having applied the two filters, we are left with a read count matrix of 16,364 genes and 1,656 cells.

##### mm10:10x dataset

(Official 10x Genomics Support, 2017). This dataset contains UMI counts. The raw UMI matrix (only the mouse part) consisted of 27,998 genes in 3,427 cells. To filter out cells with only a few expressed genes that could, for example, be generated by empty droplets, we applied a cell filter that only selected cells that expressed more than 1,500 genes. The gene filter is slightly less strict than the one for the first dataset as UMI count matrices show smaller entries (by definition several read counts collapse to less UMI counts). Thus, we filtered for genes that were expressed in at minimum ten cells with at minimum three UMIs.

#### ESTIMATION OF ONE-POPULATION MODELS

We investigated which characteristics led to the same choice of distribution for the gene expression profiles in the mm10:10x dataset. To that end, we estimated one-population models of the Poisson, NB and PB distributions for all genes and chose the most appropriate model among those three based on BIC and GOF. In Supplementary Figure S5, we visualize the values of the parameter estimates for each model and indicate the chosen models by different colors. For example (see Supplementary Figure S5B), if the NB distribution is estimated, we observe the following pattern: If the NB distribution is also the chosen one, the corresponding estimated parameters cover wide ranges *p* ∈ (0, 1) and *r* ∈ [0, 12]. In contrast, gene profiles that are most adequately described by a Poisson distribution would have resulted in a fairly large value of the parameter *p* in the NB distribution (i. e. *p* > 0.2, but more than 90% of them show *p* > 0.6) and larger values of *r* (i. e. *r* ∈ [0, 16]). Those genes that chose the PB distribution would have had smaller values in both parameters, namely *p* < 0.6 and *r* < 7.

#### BLOOD DIFFERENTIATION MARKER GENES

In Figure 3A, we observed a relatively large number of genes for which mRNA count data from Nestorowa et al. (2016) was best described by a mixture of two NB distributions rather than a zero-inflated NB distribution. In Supplementary Figure S6, we exemplarily display the count frequencies for five known blood differentiation genes from this dataset (see Paul et al., 2015), where the chosen distribution was a mixture of two NB distributions. The histograms show that some expression profiles contain many non-zero but low counts next to several large counts. Supplementary Table S3 lists the BIC values for all twelve considered models for these five genes.

#### GO TERMS

In Figure 3B, we observed a relatively large number of genes (in comparison to Figure 3A) for which mRNA count data from the mm10:10x dataset (Official 10x Genomics Support, 2017) was best described by some variant of the Poisson distribution, a distribution model that—for general contexts—is considered too simple. We thus searched for patterns in the gene ontology (GO) terms of these genes (Supplementary Figure S7.) but did not observe any apparent differences in the characteristics of the Poisson genes (i. e., those genes where the Poisson distribution was chosen) and the non-Poisson genes. To conduct this analysis, we used GO term information from http://supfam.org/SUPERFAMILY/cgi-bin/go.cgi and the R packages **biomaRt** and **GOfuncR. biomaRt** determines all GO terms of a gene, and **GOfuncR** determines all parents of a GO term. This information was then filtered for the first children GO terms.

#### OVERVIEW OF SINGLE-CELL ANALYSIS TOOLS

Many tools exist that are frequently used in single-cell analysis. In Supplementary Table S1, we provide an overview of those tools that use an underlying probability distribution to describe the counts of a specific gene’s mRNA. Most of the tools can be found at https://www.scrna-tools.org and at https://omictools.com. In the following, we describe the single categories, taken from www.scrna-tools.org. Additionally, we added the category *batch correction*.

- Batch Correction: Dealing with data from different batches
- Clustering: Unsupervised grouping of cells based on expression profiles
- Differential Expression: Testing of differential expression across groups of cells
- Dimensionality Reduction: Projection of cells into a lower-dimensional space
- Expression Patterns: Detection of genes that change over a trajectory
- Gene Networks: Identification of co-regulated gene networks
- Gene Sets: Testing or other uses of annotated gene sets
- Imputation: Estimation of expression where zeros have been observed
- Normalization: Removal of unwanted variation that may affect results
- Ordering: Ordering of cells along a trajectory
- Quality Control: Removal of low-quality cells
- Simulation: Generation of synthetic scRNA-seq datasets
- Variable Genes: Identification or use of highly (or lowly) variable genes
- Visualization: Functions for visualizing some aspect of scRNA-seq data or analysis

## DATA AND SOFTWARE AVAILABILITY

### Case Study: Simulated data

In the Case Study, we generated *in silico* data from the considered mechanistic models. For the rate sizes in the switching model, we oriented ourselves on the experimentally derived rates of Suter et al. (2011). From these, we calculated ranges for the basic and the bursting models to make simulated data comparable among models: *r*_*tran*_ = *r*_*on*_ *∪ r*_*act*_ (this is informal notation for the union of the two ranges of *r*_*on*_ and *r*_*act*_*), s*_*burst*_ = *r*_*on*_ */r*_*deact*_ and *r*_*burst*_ = *r*_*act*_. For each considered model, we generated a grid of 1,000 unique parameter sets and generated one dataset for each parameter set. The employed ranges for the parameter grid are described in Supplementary Table S4. The simulated data can be found in the GitHub repository https://github.com/fuchslab/A_mechanistic_model_for_the_negative_binomial_distribution_of_single-cell_mRNA_counts.

### Scripts

All scripts used in this study can be found in our open GitHub repository https://github.com/fuchslab/A_mechanistic_model_for_the_negative_binomial_distribution_of_single-cell_mRNA_counts.

### Software

The newly generated R package **scModels** can be found on CRAN and in our open GitHub repository https://github.com/fuchslab/scModels.

## SUPPLEMENTARY MATERIALS

### SINGLE CELL ANALYSIS TOOLS

In Supplementary Table S1, an overview of tools in single-cell analysis is given that are based on distributional assumptions.

**Table S1:** Related to Appendix. Overview of single-cell analysis tools with underlying distributional assumptions. In italics we highlight those categories that were assigned by ourselves to tools that were not listed on www.scrna-tools.org.

| Tool | Category | NB | PB | Other | ZI | Hurdle | Notes |
| --- | --- | --- | --- | --- | --- | --- | --- |
| BASICS | Normalization, Differential Expression, Variable Genes, Simulation | x |  |  |  |  | Poisson-gamma, Bayesian hierarchical models, Vallejos et al. (2015) |
| bayNorm | Normalization, Imputation, Simulation | x |  |  |  |  | Binomial sampling with NB priors, Tang et al. (2018) |
| BEAM | *Ordering, Expression Patterns, Differential Expression* | x |  |  |  |  | Branch-dependent gene expression as a contrast between two NB GLMs, Qiu et al. (2017) |
| BPSC | Differential Expression |  | x |  | x |  | BP3 is PB; BP4 adds fractions scaling parameter; BP5 adds ZI; Vu et al. (2016) |
| ComBat | *Batch Correction* |  |  | x |  |  | Uses normal distribution on normalized data, Stein et al. (2015) |
| DCA | Imputation | x |  |  | x |  | Eraslan et al. (2019) |
| DPT | Ordering, Expression Patterns, Visualization |  |  | x |  | x | Normal distribution, Haghverdi et al. (2016) |
| D3E | Differential Expression |  | x |  |  |  | Delmans and Hemberg (2016) |
| diffxPy | *Differential Expression* | x |  |  | x |  | https://github.com/theislab/diffxpy |
| limma | *Normalization, Differential Expression, Gene Sets, Batch Correction* |  |  | x |  |  | Linear model using normal distributions, Ritchie et al. (2015) |
| lineagePulse | Differential Expression, Expression Patterns, Visualization, Simulation | x |  |  | x |  | https://github.com/YosefLab/LineagePulse |
| MAST | Quality Control, Normalization, Differential Expression, Gene Sets, Gene Networks |  |  |  |  | x | Logistic regression & Gaussian linear model for expressed genes, Finak et al. (2015) |
| M3Drop | Differential Expression, Marker Genes, Visualization, Simulation | x |  |  |  |  | Depth-adjusted NB, Andrews and Hemberg (2018) |
| powSimR | Visualization, Simulation | x |  |  | x |  | The user has the option to include zero inflation (default is not to use it), Vieth et al. (2017) |
| SAVER | Imputation | x |  |  |  |  | Huang et al. (2018) |
| SCDE | Differential Expression, Gene Sets, Visualization |  |  | x | (x) |  | Poisson-NB mixture: Poisson for dropout, NB for amplified expression, Kharchenko et al. (2014) |
| SCHiRM | Normalization, Gene Networks, Visualization, Simulation |  |  | x |  |  | Poisson-lognormal, Intosalmi et al. (2018) |
| scImpute | Imputation |  |  | x | (x) |  | Gamma-normal mixture on log-transformed expression: dropouts modeled via normal distribution, Li and Li (2018) |
| sctransform | Normalization, Integration, Differential Expression, Transformation, Visualization | x |  |  |  |  | Regularized NB regression, Hafemeister and Satija (2019) |
| scVI | Dimensionality Reduction | x |  |  | x |  | ZINB-like generative model, Lopez et al. (2018) |
| Splatter | Visualization, Simulation | x |  |  | x |  | Some intermediate steps; gene- and cell-wise mean are modeled with gamma distribution, Zappia et al. (2017) |
| ZIFA | Dimensionality Reduction |  |  | x | x |  | Zero-inflated Gaussian (Bernoulli-normal mixture), Pierson and Yau (2015) |
| ZINB-WaVE | Normalization, Dimensionality Reduction, Simulation | x |  |  | x |  | Risso et al. (2018) |

### SWITCHING MODEL

In Supplementary Figure S1, we depicted an alternative description of the switching process shown in Figure 1B that is required for the calculations in Appendix.

**Figure S1:**
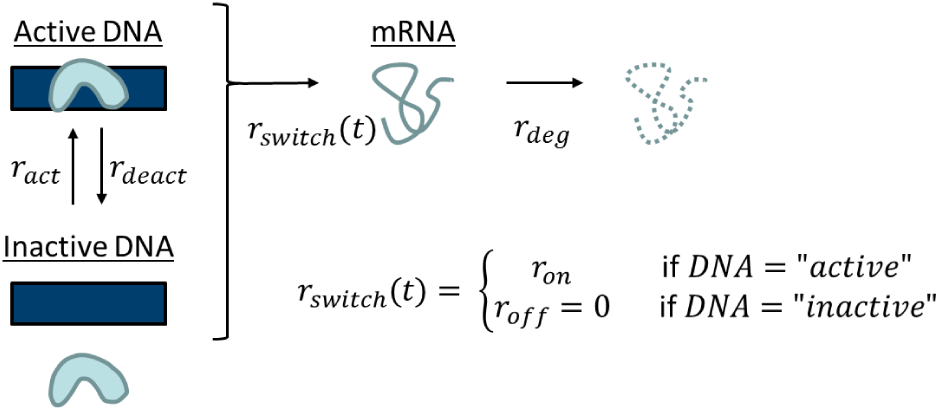
Related to Figure 1B and to Appendix. Detailed depiction of the Markov chain that governs the switching process in the switching model of gene activation, transcription and degradation.

### BURSTING MODEL

In Supplementary Figure S2, all possible state transitions of the bursting model are depicted. This is the basis for deriving the master equation of the model as shown in the Appendix.

**Figure S2:**
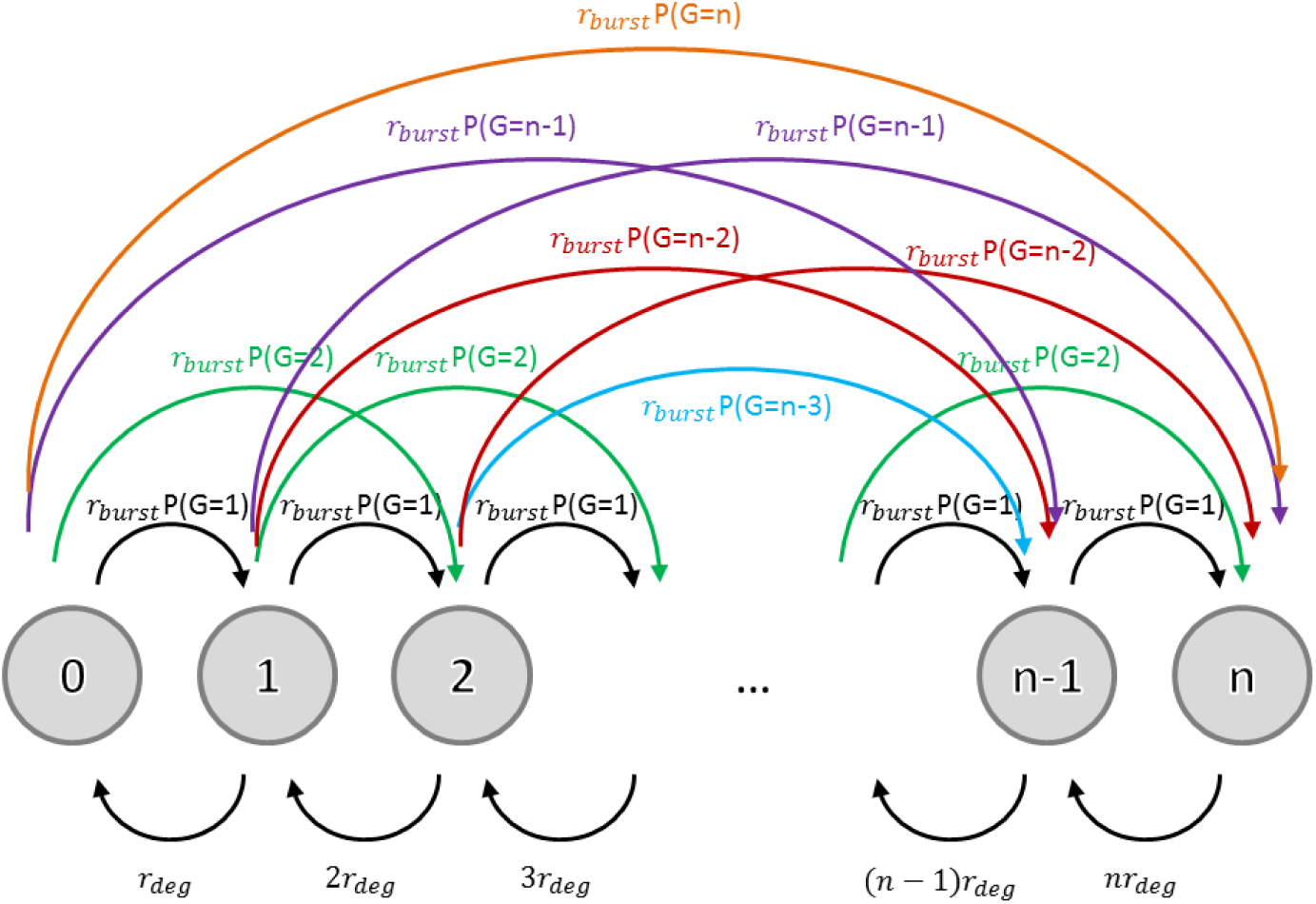
Related to Figure 1D and to Appendix. Bursting model with all states and possible transitions between states, assuming that at most one event (transcription or degradation) can happen at the same time. Transitions from one node to itself are not depicted. Here, *P* (*G* = *k*) stands for the probability of a geometrically distributed random variable *G* taking the value *k*.

### KUMMER FUNCTION AND scModels

Supplementary Figure S3 shows the incomplete implementation of the Kummer function that is contained in **dAsianOptions** and our fix that is described in more detail in the Appendix.

**Figure S3:**
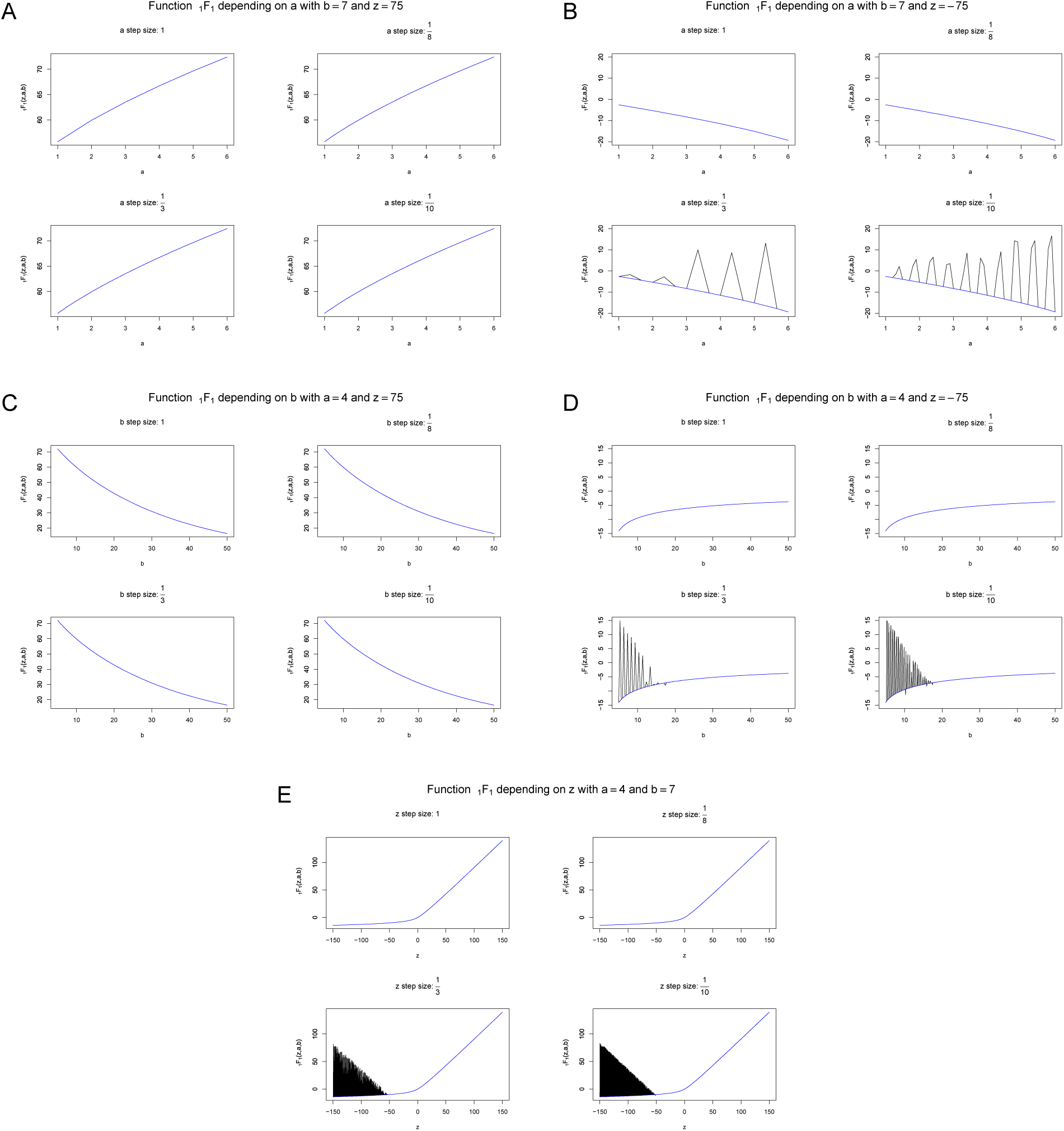
Related to Appendix. Behavior of the Kummer function for different parameter sets based on the implementations of **dAsianOptions** in black and **scModels** in blue. (A,C and E) As long as *z* is positive, the Kummer function of both packages return the correct values. (B,D and E) As soon as *z* is negative (smaller than −50) the Kummer function of the **fAsianOptions** returns wrong values for *a, b* and *z* values that cannot be expressed by the general formula *m* 2^*-n*^, *m, n*∈ ℕ_0_. This bug is fixed in the new implementation of the Kummer function in **scModels**.

Supplementary Table S2 shows the results of the simulation study on package comparisons that is explained in the Appendix.

**Table S2:**
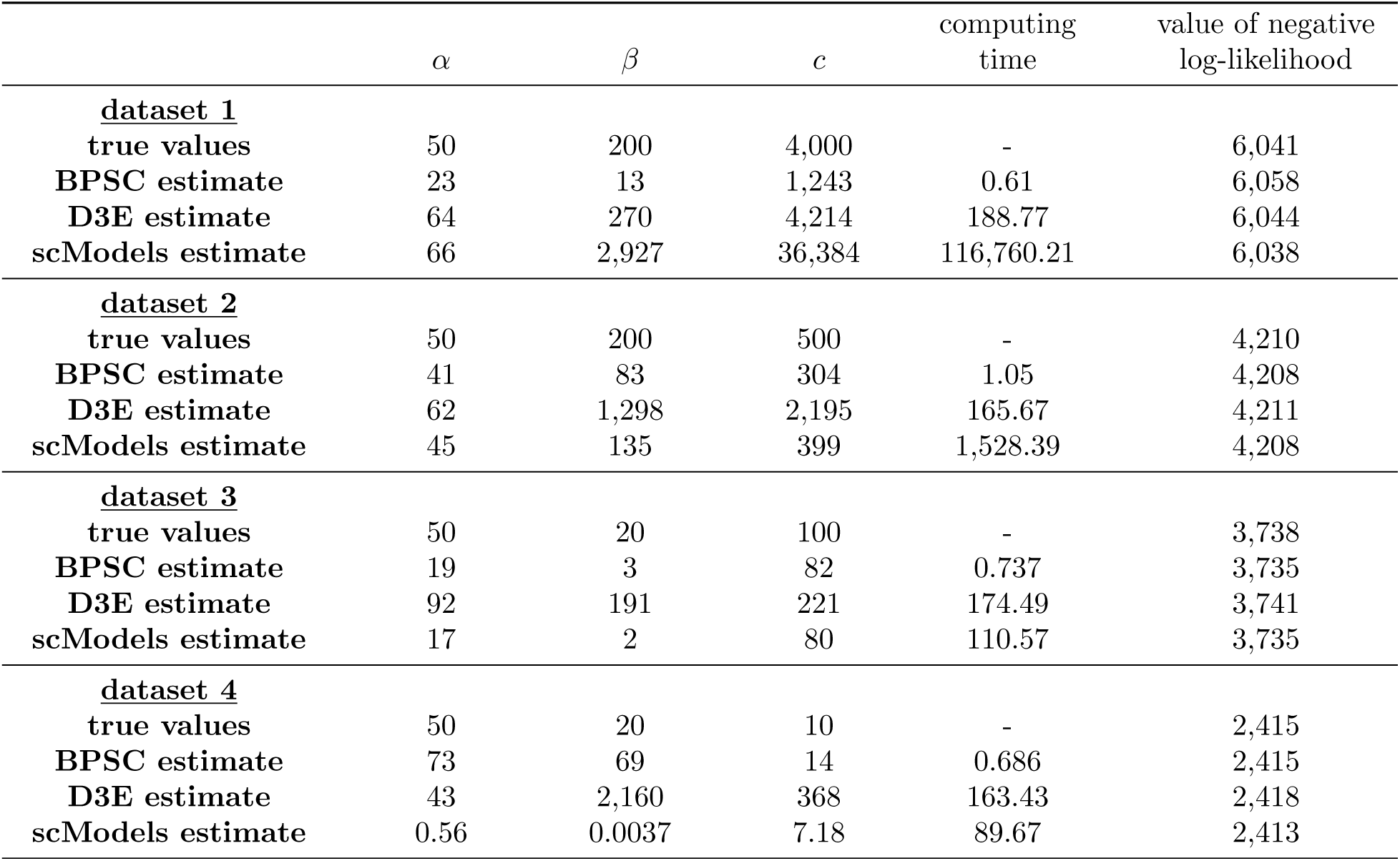
Related to Appendix. Results of parameter estimation for the Poisson-beta distribution using the software packages **BPSC, D3E** and **scModels**. We simulated four datasets of size 1,000 each (for details, see Appendix). The table shows values of the parameters *α, β* and *c*: the true values used for synthetic data generation, and the estimates obtained through application of the different packages. The last two columns show the computation time measured in seconds for each algorithm and the value of the negative log-likelihood function (computed using the function scModels::nlogL pb()) evaluated at the respective parameter values. Smaller values of the negative log-likelihood indicate better point estimates.

| | $\alpha$ | $\beta$ | $c$ | computing<br>time | value of negative<br>log-likelihood |
| --- | --- | --- | --- | --- | --- |
| **<u>dataset 1</u>** |  |  |  |  |  |
| true values | 50 | 200 | 4,000 | - | 6,041 |
| BPSC estimate | 23 | 13 | 1,243 | 0.61 | 6,058 |
| D3E estimate | 64 | 270 | 4,214 | 188.77 | 6,044 |
| scModels estimate | 66 | 2,927 | 36,384 | 116,760.21 | 6,038 |
| **<u>dataset 2</u>** |  |  |  |  |  |
| true values | 50 | 200 | 500 | - | 4,210 |
| BPSC estimate | 41 | 83 | 304 | 1.05 | 4,208 |
| D3E estimate | 62 | 1,298 | 2,195 | 165.67 | 4,211 |
| scModels estimate | 45 | 135 | 399 | 1,528.39 | 4,208 |
| **<u>dataset 3</u>** |  |  |  |  |  |
| true values | 50 | 20 | 100 | - | 3,738 |
| BPSC estimate | 19 | 3 | 82 | 0.737 | 3,735 |
| D3E estimate | 92 | 191 | 221 | 174.49 | 3,741 |
| scModels estimate | 17 | 2 | 80 | 110.57 | 3,735 |
| **<u>dataset 4</u>** |  |  |  |  |  |
| true values | 50 | 20 | 10 | - | 2,415 |
| BPSC estimate | 73 | 69 | 14 | 0.686 | 2,415 |
| D3E estimate | 43 | 2,160 | 368 | 163.43 | 2,418 |
| scModels estimate | 0.56 | 0.0037 | 7.18 | 89.67 | 2,413 |

Supplementary Figure S4 shows further results from the package comparison simulation study.

**Figure S4:**
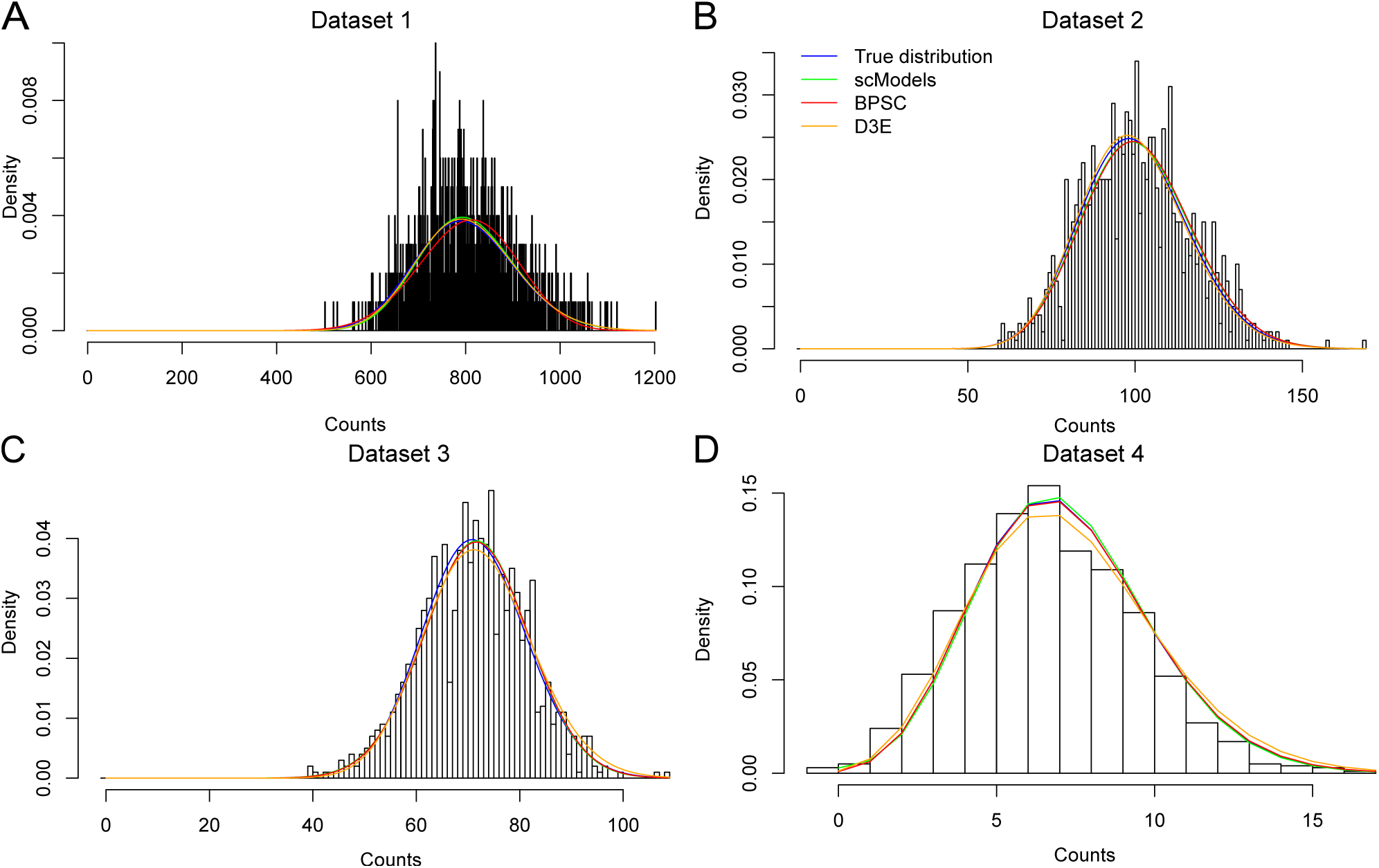
Related to Appendix and Table S2. Histograms of the four simulated datasets (A-D) and Poisson-beta densities using the true and estimated parameters from Table S2, respectively: true (blue), **scModels** (green), **BPSC** (red) and **D3E** (orange).

### ESTIMATION OF ONE-POPULATION MODELS

Supplementary Figure S5 shows the results when forcing each gene of the mm10:10x dataset to be modeled by a one-population distribution.

**Figure S5:**
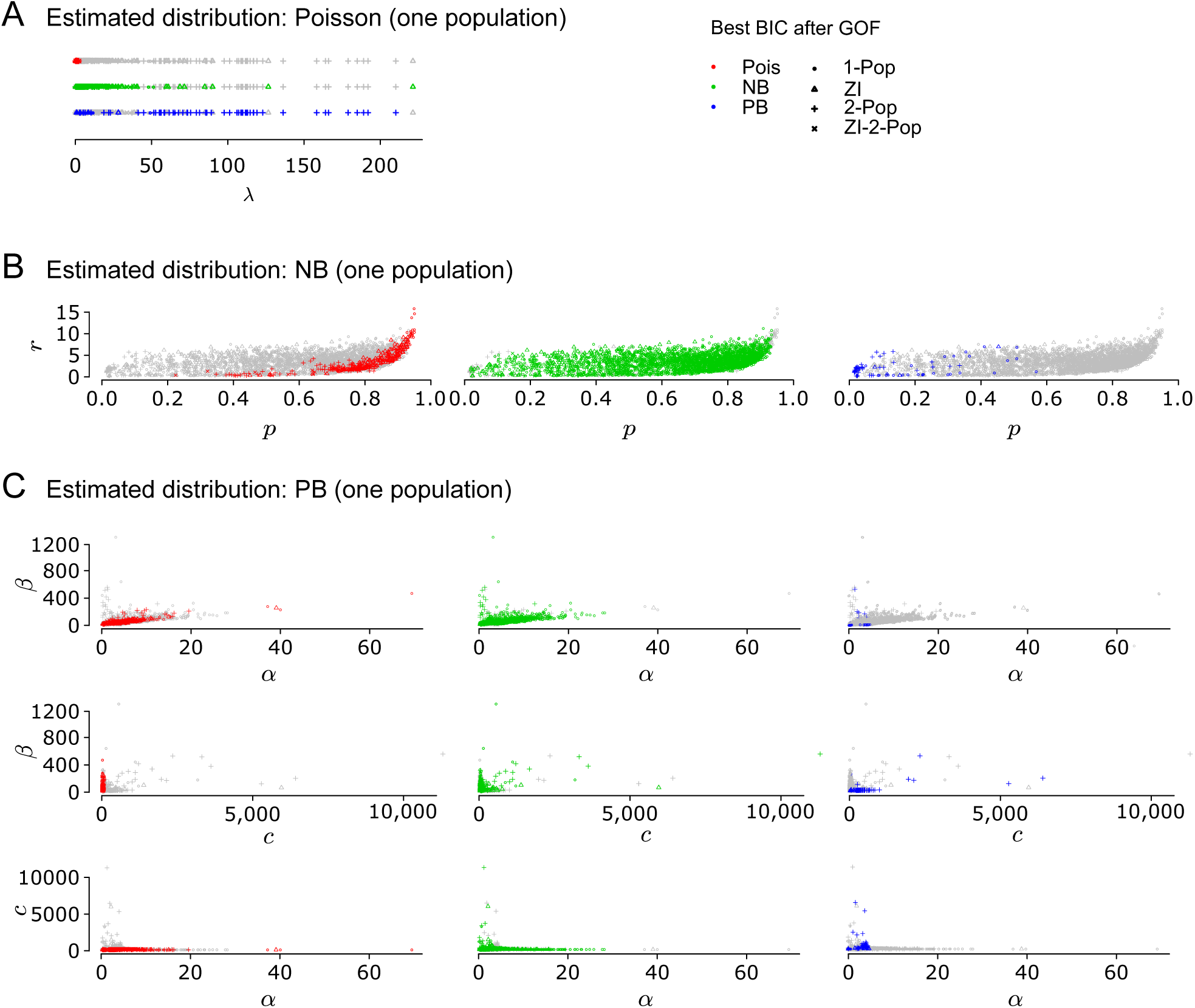
Related to Figure 3B. We estimated one-population models of the Poisson, NB and PB distributions for all genes in the mm10:10x dataset and chose the most appropriate model based on BIC after GOF. (A) Estimated *λ* parameters for the Poisson distribution. Each dot corresponds to one gene. In the top line, estimated values are coloured in red for those genes where the Poisson distribution was chosen. In the middle line, green symbols indicate estimated values in the Poisson model where the NB distribution would have been preferred. In the bottom line, blue colour indicates the estimates for those genes that chose the PB distribution. (B) Similarly for the NB distribution. (C) Similarly for the PB distribution.

### BLOOD DIFFERENTIATION MARKER GENES

In Supplementary Figure S6 we plotted for exemplary reasons some known blood differentiation genes (see Paul et al., 2015).

**Figure S6:**
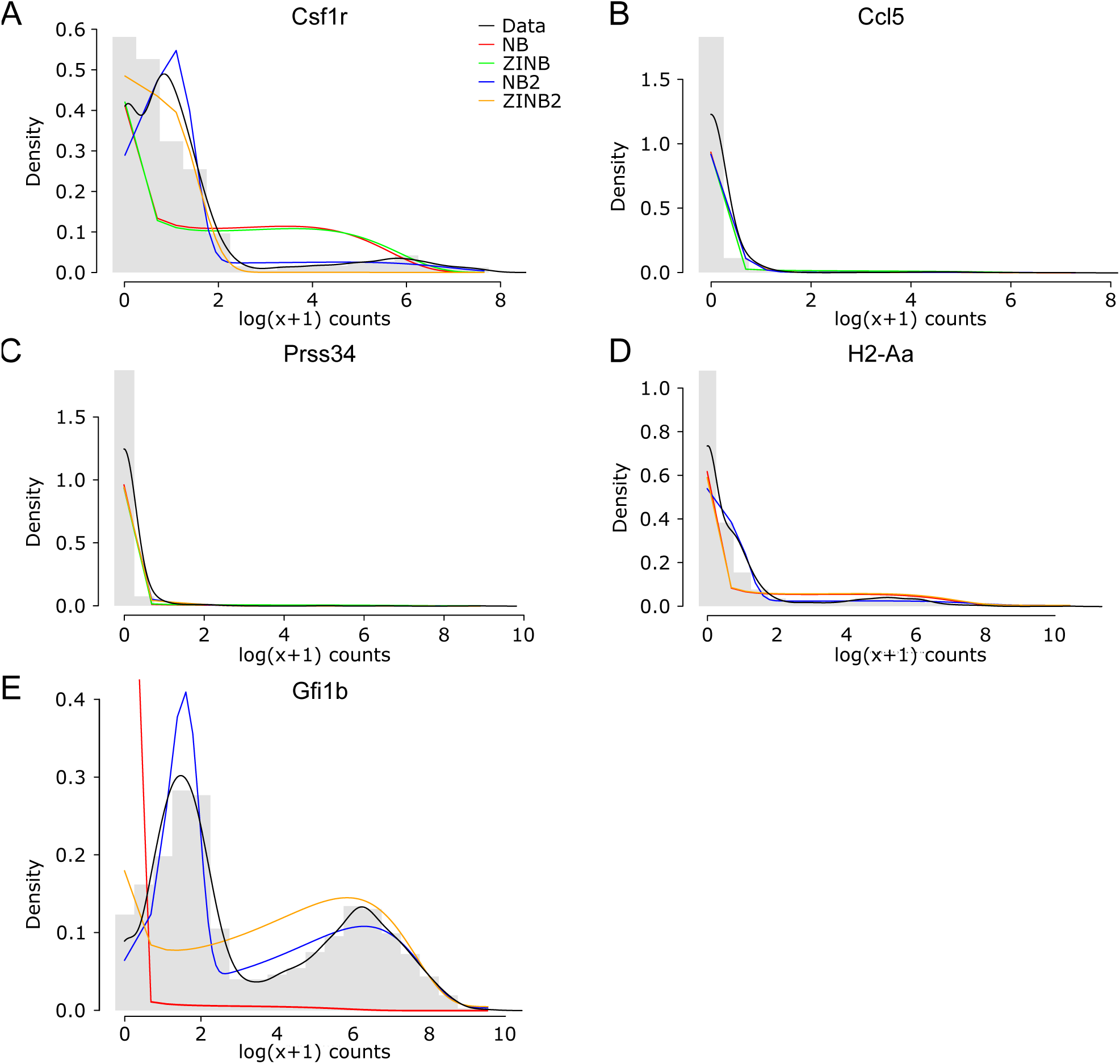
Related to Figure 3A. (A)-(E) Log-transformed mRNA count profiles for five genes (based on 1,656 single cells) from the dataset by Nestorowa et al. (2016), known as blood differentiation markers. Coloured lines indicate the densities of the estimated empirical and NB distribution variants: empirical distribution (data, black), NB distribution (NB, red), zero-inflated NB distribution (ZINB, green), mixture of two NB distributions (NB2, blue), zero-inflated mixture of two NB distributions (ZINB2, yellow). The blue NB2 was the most appropriate distribution in all cases.

Supplementary Table S3 contains more estimation details for these five genes.

**Table S3:** Related to Figures 3A and S6 and Appendix. BIC values for selected blood differentiation marker genes (based on 1,656 single cells) as described in Supplementary Figure S6 and the text body of the Supplementary Materials. *Columns:* Results for five genes Csf1r, Ccl5, Prss34, H2-Aa, Gfi1b. *Rows:* BIC values for all twelve estimated models; the selected model and the corresponding p-value of the GOF test (for all gene profiles, the NB2 model is chosen); percentages of zero counts, one counts, and counts larger than one.

|  | Csf1r | Ccl5 | Prss34 | H2-Aa | Gfi1b |
| --- | --- | --- | --- | --- | --- |
| BIC<sub>Pois</sub> (1 parameter) | 377,837 | 62,721 | 194,047 | 1,224,382 | 1,690,010 |
| BIC<sub>ZIPois</sub> (2 parameters) | 330,413 | 29,741 | 107,502 | 984,306 | 1,618,095 |
| BIC<sub>Pois2</sub> (3 parameters) | 60,829 | 9,930 | 20,889 | 390,636 | 489,696 |
| BIC<sub>ZIPois2</sub> (4 parameters) | 74,845 | 8,759 | 150,63 | 568,092 | 502,270 |
| BIC<sub>NB</sub> (2 parameters) | 10,295 | 1,878 | 1,545 | 8,454 | 29,842 |
| BIC<sub>ZINB</sub> (3 parameters) | 10,292 | 1,865 | 1,490 | 8,454 | 18,653 |
| BIC<sub>NB2</sub> (5 parameters) | 8,505 | 1,693 | 1,387 | 7,672 | 17,585 |
| BIC<sub>ZINB2</sub> (6 parameters) | 9,978 | 1,713 | 1,401 | 8,477 | 18,676 |
| BIC<sub>PB</sub> (3 parameters) | 10,407 | 1,865 | 1,490 | 8,516 | 18,675 |
| BIC<sub>ZIPB</sub> (4 parameters) | 10,414 | 1,897 | 1,555 | 8,618 | 18,803 |
| BIC<sub>PB2</sub> (7 parameters) | 10,187 | 1,920 | 1,570 | 8,467 | 18,778 |
| BIC<sub>ZIPB2</sub> (8 parameters) | 10,727 | 1,937 | 1,653 | 8,564 | 18,787 |
| Selected model | NB2 | NB2 | NB2 | NB2 | NB2 |
| p-value of GOF ($\chi^2$) test | 1.515e-02 | 9.577e-01 | 1.054e-05 | 9.371e-01 | 2.233e-04 |
| Percentage of zero counts | 29.0% | 91.4% | 93.5% | 54.0% | 6.2% |
| Percentage of one counts | 26.3% | 5.6% | 3.7% | 19.1% | 8.1% |
| Percentage of counts larger than one | 44.6% | 3.0% | 2.7% | 26.9% | 85.7% |

### GO TERMS

Supplementary Figure S7 shows a GO term analysis comparing groups of genes which where best described by a Poissonian model and those that were not.

**Figure S7:**
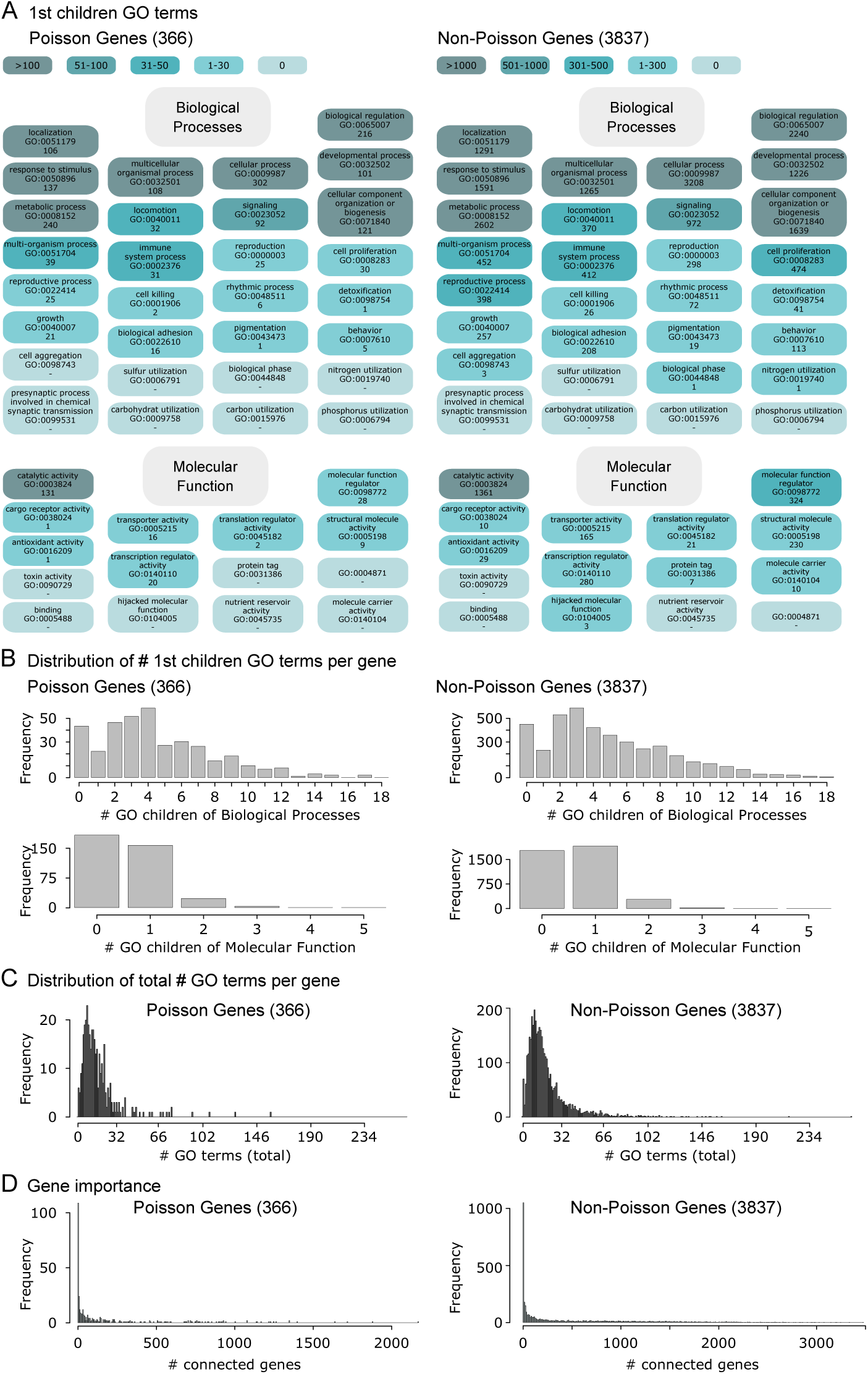
Related to Figure 3B. (A) Amount of Poisson and non-Poisson genes (after GOF) that are contained in the first level of GO term children of the families *biological process* and *molecular function*. (B) Distribution of the first children GO terms of the families *biological process* and *molecular function* for Poisson and non-Poisson genes. (C) Distribution of the overall number of GO terms a gene is contained in. GO terms were taken from the initial **biomaRt** determination. (D) Gene importance of Poisson and non-Poisson genes: Functional coupling network of genes taken from funcoup.sbc.su.se. Each link with weight > 0.75 was taken and the distribution of the number of coupled genes per gene in this network is plotted.

### Case Study

Supplementary Table S4 shows the ranges of the rates used in the simulation study in the Case Study.

**Table S4:** Related to Figure 4. Ranges of rates in the simulation study in the Case Study.

| **Mechanistic model** | **Rate parameter** | **Minimum value** | **Maximum value** |
| --- | --- | --- | --- |
| Basic model | $r_{tran}$ | 0.005 | 2.5 |
| | $r_{deg}$ | 0.001 | 0.05 |
| Bursting model | $r_{burst}$ | 0.005 | 0.06 |
| | $s_{burst}$ | 0.5 | 2.5 |
| | $r_{deg}$ | 0.001 | 0.05 |
| Switching model | $r_{act}$ | 0.005 | 0.06 |
| | $r_{deact}$ | 0.01 | 0.6 |
| | $r_{on}$ | 0.5 | 2.5 |
| | $r_{deg}$ | 0.001 | 0.05 |

